## Supplementary material for "Profiling cell envelope-antibiotic interactions reveals vulnerabilities to β-lactams in a multidrug-resistant bacterium": File for all Supplemental Figures and Tables

**Supporting Tables S1 – S10** are given here

**Supporting Figures S1 – S13** are given here

**Supporting Data S1** is provided as a separate Excel file

**Table S1** Summary statistics of the randomly barcoded transposon mutant library

| Category | K56-2 Library |
|---|---|
| Number of mutant colonies recovered | ~815, 000 |
| Number of unique barcodes | 338,116 |
| Number of unique insertion positions | 145,585 |
| Protein-coding genes with (or without) central insertions (10-90% of gene) | 6412 (561) |
| Mean/median strains per interrupted protein-coding gene | 31.9 / 12 |
| Gene and transposon on same strand | 49.9% |
| Average distance between insertions | 18.7 bp |

**Table S2.** Effect of CRISPRi knockdown of β-lactamase genes on susceptibility to AMP, CFD, TAZ, and CTT. MIC values are the median of three replicates and are reported in µg/mL. Bolded values indicate at least 2-fold change.

| Antibiotic | Gene | MIC (µg/mL) -Rha | +Rha | Fold MIC* |
|---|---|---|---|---|
| AMP | NTC | >512 | >512 | - |
| | *penR* | >512 | >512 | - |
| | *bla*$_{AmpC}$ | >512 | >512 | - |
| | *bla*$_{PenB}$ | >512 | **256** | **>2** |
| CFD | NTC | 0.125 | 0.125 | 1 |
| | *penR* | 0.125 | 0.125 | 1 |
| | *bla*$_{AmpC}$ | 0.125 | 0.125 | 1 |
| | *bla*$_{PenB}$ | 0.125 | 0.125 | 1 |
| TAZ | NTC | 128 | 128 | 1 |
| | *penR* | 128 | **64** | **2** |
| | *bla*$_{AmpC}$ | 128 | 128 | **1** |
| | *bla*$_{PenB}$ | 128 | **32** | **4** |
| CTT | NTC | 64 | 64 | 1 |
| | *penR* | 64 | **16** | **4** |
| | *bla*$_{AmpC}$ | 64 | 64 | 1 |
| | *bla*$_{PenB}$ | 64 | 64 | 1 |

*Fold MIC is the ratio of the MIC -Rha to the MIC +Rha

**Table S3.** Susceptibility of Gram-negative panel to AVI combinations (± 8 µg/mL AVI). MICs are the median of three replicates and are reported in µg/mL.

| Species | Strain | AZT | AVI/AZT | Ratio | Mero | AVI/MEM | Ratio | CAZ | AVI/CAZ | Ratio | CFD | AVI/CFD | Ratio |
|---|---|---|---|---|---|---|---|---|---|---|---|---|---|
| *A. xylosoxidans* | ACH02 | 64 | 64 | 1 | 1 | 1 | 1 | 2 | 2 | 1 | 0.25 | 0.25 | 1 |
| | ACH03 | 32 | 32 | 1 | 0.25 | 0.25 | 1 | 1 | 1 | 1 | 0.125 | 0.125 | 1 |
| | ACH09 | 16 | 16 | 1 | 0.125 | 0.125 | 1 | 1 | 1 | 1 | 0.25 | 0.25 | 1 |
| *S. maltophilia* | D457 | 16 | 2 | **8** | 32 | 32 | 1 | 1 | 1 | 1 | 0.125 | 0.125 | 1 |
| | K279a | 128 | 1 | **128** | 2 | 2 | 1 | 2 | 1 | **2** | 0.125 | 0.125 | 1 |
| | 03 T2510 | 256 | 4 | **64** | >64 | >64 | - | 32 | 32 | 1 | 1 | 1 | 1 |
| | 04 T22674 | 64 | 1 | **64** | 16 | 16 | 1 | 2 | 2 | 1 | 0.5 | 0.5 | 1 |
| | 05 41846 | 128 | 2 | **64** | >64 | >64 | - | 4 | 4 | 1 | 0.5 | 0.5 | 1 |
| | 21NE928511 | 256 | 4 | **64** | >64 | >64 | - | >128 | >128 | **-** | >4 | >4 | - |
| | 20NC580771 | 256 | 2 | **128** | >64 | >64 | - | >128 | >128 | **-** | 0.25 | 0.25 | 1 |
| *P. aeruginosa* | PAO1 | 4 | 4 | 1 | 0.5 | 0.5 | 1 | 1 | 1 | 1 | 1 | 1 | 1 |
| | PA14 | 8 | 8 | 1 | 0.5 | 0.25 | **2** | 2 | 2 | 1 | 1 | 1 | 1 |
| | 70 | 0.25 | 0.25 | 1 | 0.125 | 0.125 | 1 | 0.5 | 0.5 | 1 | 0.5 | 0.5 | 1 |
| | 71 | 128 | 8 | **16** | 32 | 4 | **8** | >128 | 4 | **>32** | 4 | 4 | 1 |
| | 74 | 128 | 2 | **64** | 16 | 2 | **8** | 128 | 2 | **64** | >4 | >4 | - |
| | 20NB681421 | 16 | 16 | 1 | 1 | 0.5 | **2** | 4 | 4 | 1 | 1 | 1 | 1 |
| | 21NE72623 | 4 | 2 | **2** | 0.5 | 0.125 | **4** | 16 | 2 | **8** | >4 | >4 | - |
| *B. cenocepacia* | 110041 | 64 | 16 | **4** | 32 | 4 | **8** | 4 | 4 | 1 | 0.0625 | 0.0625 | 1 |
| | BC7 | >128 | >128 | - | 16 | 1 | **16** | 16 | 16 | 1 | >4 | >4 | - |
| | K56-2 | 256 | 8 | **32** | 16 | 2 | **8** | 32 | 4 | **8** | 0.125 | 0.125 | 1 |
| | VC10414 | 16 | 8 | **2** | 32 | 2 | **16** | 8 | 4 | **2** | 0.125 | 0.125 | 1 |
| | VC14488 | 16 | 2 | **8** | 4 | 1 | **4** | 4 | 2 | **2** | 0.25 | 0.25 | 1 |
| | VC14543 | 256 | 32 | **8** | 16 | 4 | **4** | 8 | 8 | 1 | 0.125 | 0.0625 | **2** |
| | VC14761 | 8 | 2 | **4** | 4 | 1 | **4** | 2 | 2 | 1 | 0.125 | 0.0625 | **2** |
| | F4200989-3 | >256 | 64 | **>4** | 32 | 4 | **8** | 128 | 32 | **4** | 2 | 1 | **2** |
| | G11815681 | >256 | 32 | **>8** | 32 | 2 | **16** | 16 | 8 | **2** | 2 | 0.5 | **4** |
| | F3071909 | 256 | 32 | **8** | 16 | 2 | **8** | 32 | 32 | 1 | >4 | >4 | - |
| | F9141525 | >256 | 32 | **>8** | 32 | 4 | **8** | 128 | 16 | **8** | 0.5 | 0.5 | 1 |
| | 130723 | 256 | 8 | **32** | 4 | 1 | **4** | 16 | 8 | **2** | 0.5 | 0.25 | 2 |

| | | | | | | | | | | | | | |
|---|---|---|---|---|---|---|---|---|---|---|---|---|---|
| | 134991 | 16 | 16 | 1 | 2 | 2 | 1 | 8 | 8 | 1 | >4 | >4 | - |
| *B. contaminans* | LMG23361 | 32 | 2 | **16** | 4 | 0.5 | **8** | 2 | 2 | 1 | 0.125 | 0.125 | 1 |
| | 21NG609698 | 256 | 8 | **32** | 32 | 8 | **4** | 16 | 4 | **4** | 2 | 2 | 1 |
| | 21NG595759 | 256 | 8 | **32** | 32 | 8 | **4** | 16 | 8 | **2** | 1 | 2 | 0.5 |
| *B. dolosa* | CEP021 | 8 | 8 | 1 | 2 | 1 | **2** | 2 | 1 | **2** | 0.5 | 0.5 | 1 |
| | AU0645 | >256 | 16 | **>16** | 16 | 2 | **8** | >128 | 8 | **>16** | >4 | >4 | - |
| | LM018941 | >256 | 4 | **>64** | 16 | 0.5 | **32** | 16 | 2 | **8** | >4 | >4 | - |
| | AU2167 | >256 | 16 | **>16** | 32 | 1 | **32** | 64 | 4 | **16** | >4 | >4 | - |
| | AU6940 | >256 | 8 | **>32** | 32 | 2 | **16** | 16 | 16 | 1 | 2 | 2 | 1 |
| *B. gladioli* | VC14812 | 128 | 32 | **4** | 8 | 8 | 1 | 32 | 16 | **2** | 0.5 | 0.5 | 1 |
| | VC19233 | 32 | 8 | **4** | 1 | 1 | 1 | 4 | 2 | **2** | 0.25 | 0.25 | 1 |
| | 113611 | 128 | 32 | **4** | 2 | 2 | 1 | 32 | 32 | 1 | 4 | 4 | 1 |
| | 132208 | 4 | 4 | 1 | 0.5 | 0.5 | 1 | 2 | 2 | 1 | 0.5 | 0.5 | 1 |
| *B. multivorans* | 130034 | 16 | 8 | **2** | 8 | 4 | **2** | 8 | 8 | 1 | 0.125 | 0.125 | 1 |
| | C5393 | 64 | 8 | **8** | 16 | 1 | **16** | 8 | 4 | **2** | 0.25 | 0.25 | 1 |
| | VC15535 | 256 | 16 | **16** | 16 | 8 | **2** | 32 | 16 | **2** | 0.5 | 0.5 | 1 |
| | VC15555 | 4 | 4 | 1 | 8 | 4 | **2** | 2 | 2 | 1 | >4 | >4 | - |
| | VC19694 | 128 | 4 | **32** | 8 | 4 | **2** | 4 | 1 | **4** | 1 | 1 | 1 |
| | VC9825 | 8 | 4 | **2** | 2 | 1 | **2** | 2 | 2 | 1 | 0.25 | 0.125 | **2** |
| | 131077 | 16 | 8 | **2** | 4 | 1 | **4** | 8 | 8 | 1 | 0.125 | 0.125 | 1 |
| | 134955 | 16 | 4 | **4** | 8 | 1 | **8** | 8 | 8 | 1 | 0.125 | 0.125 | 1 |
| | F9091687 | 2 | 0.5 | **4** | 0.5 | 0.5 | 1 | 2 | 2 | 1 | 0.25 | 0.25 | 1 |
| | LM013010 | 64 | 4 | **16** | 8 | 1 | **8** | 4 | 4 | 1 | 0.25 | 0.25 | 1 |
| | V1120157 | 4 | 4 | 1 | 1 | 1 | 1 | 2 | 2 | 1 | 0.125 | 0.125 | 1 |
| *B. vietnamiensis* | CEP40 | 4 | 2 | **2** | 1 | 1 | 1 | 4 | 2 | **2** | 0.125 | 0.0625 | **2** |
| | VC11431 | 256 | 4 | **64** | 16 | 1 | **16** | 32 | 2 | **16** | 1 | 0.125 | **8** |
| | VC18984 | 256 | 8 | **32** | 8 | 4 | **2** | 16 | 4 | **4** | >4 | >4 | - |

**Table S4.** MIC values of other β-lactams in K56-2 with AVI and TAZ. MIC values are the median of three replicates and are reported in µg/mL. Bolded values indicate changes upon β-lactamase addition.

| | **β-Lactam Alone** | **+ 8 µg/mL AVI** | **+ 8 µg/mL TAZ** |
|---|---|---|---|
| Ceftazidime | 32 | **4** | 32 |
| Cefotaxime | >128 | **16** | >128 |
| Cephalexin | >128 | **128** | >128 |
| Cefmetazole | 128 | 128 | 128 |
| Cefoperazone | >128 | **64** | >128 |
| Ceftriaxone | >128 | **4** | >128 |
| Moxalactam | 128 | 128 | 128 |

**Table S5.** Susceptibility of CF pathogens and clinical isolates to cefiderocol and ceftazidime in CAMHB and M9+CAA. MIC values (µg/mL) are medians of at least three biological replicates. The ratio is the MIC in M9+CAA divided by the MIC in CAMHB. Bold indicates at least 2-fold difference between medium types.

| | | CFD | | | CAZ | | |
|---|---|---|---|---|---|---|---|
| Species | Strain | CAMHB | M9+CAA | Ratio | CAMHB | M9+CAA | Ratio |
| *A. xylosoxidans* | ACH02 | 0.25 | 0.00781 | **0.03125** | 2 | 2 | 1 |
| | ACH03 | 0.125 | 0.03125 | **0.25** | 1 | 1 | 1 |
| *S. maltophilia* | D457 | 0.125 | 0.25 | **2** | 1 | 1 | 1 |
| | K279a | 0.125 | 0.125 | 1 | 2 | 4 | **2** |
| *P. aeruginosa* | PAO1 | 1 | 0.0078 | **0.0078** | 1 | 1 | 1 |
| | PA14 | 1 | 0.0078 | **0.0078** | 2 | 2 | 1 |
| *B. cenocepacia* | 110041 | 0.0625 | 0.5 | **8** | 4 | 2 | **0.5** |
| | BC7 | >4 | >4 | N/A | 16 | 16 | 1 |
| | K56-2 | 0.125 | 1 | **4** | 32 | 32 | 1 |
| | VC10414 | 0.125 | 0.03125 | **0.25** | 8 | 8 | 1 |
| | VC14488 | 0.25 | 4 | **16** | 4 | 8 | **2** |
| | VC14543 | 0.125 | 1 | **8** | 8 | 4 | **0.5** |
| | VC14761 | 0.125 | 0.125 | 1 | 4 | 8 | **2** |
| *B. dolosa* | CEP021 | 0.5 | >4 | **>8** | 2 | 2 | 1 |
| *B. multivorans* | 130034 | 0.125 | 0.5 | **4** | 8 | 8 | 1 |
| | C5393 | 0.25 | 0.5 | **2** | 8 | 8 | 1 |
| | VC15535 | 0.5 | >8 | **>16** | 32 | 32 | 1 |
| | VC19694 | 1 | 4 | **4** | 4 | 4 | 1 |
| | VC9825 | 0.25 | 2 | **8** | 2 | 4 | **2** |
| | ATCC17616 | 4 | 4 | 1 | 8 | 8 | 1 |
| *B. vietnamiensis* | CEP40 | 0.125 | 2 | **16** | 4 | 4 | 1 |
| | VC11431 | 1 | 4 | **4** | 32 | 32 | 1 |
| | VC18984 | >8 | >8 | N/A | 16 | 8 | **0.5** |
| *B. thailandensis* | E264 | 0.125 | 0.0625 | **0.5** | 8 | 4 | **0.5** |

**Table S6.** Whole panel of CRISPRi mutants in genes associated with iron uptake from Figure 8 and their susceptibility to cefiderocol and ceftazidime in CAMHB and M9+CAA. MIC values (µg/mL) presented are medians of at least four biological replicates in the presence of 0.5% rhamnose. Bold indicates at least 2-fold difference from the NTC. Gene names are assigned by homology, if known, or else the locus tag is given (preceded by K562_).

| Gene Function | CRISPRi Target | CFD | | CAZ | |
|---|---|---|---|---|---|
| | | CAMHB | M9+CAA | CAMHB | M9+CAA |
| N/A | NTC | 0.125 | 1 | 32 | 32 |
| TBDR (Outer membrane protein) | *piuA* (RS21350) | **0.5** | **4** | 32 | 32 |
| | RS00600 | 0.125 | 1 | 32 | 32 |
| | RS07175 | 0.125 | 1 | 32 | 32 |
| | RS09770 | 0.125 | 1 | 32 | 32 |
| | RS09805 | 0.125 | 1 | 32 | 32 |
| | *cntO* (RS10150)* | **0.25** | **2** | **8** | **16** |
| | *orbA* (RS10200) | 0.125 | 1 | 32 | 32 |
| | *fecA* (RS11910) | 0.125 | 1 | 32 | 32 |
| | RS12040 | 0.125 | 1 | 32 | 32 |
| | RS14405 | 0.125 | 1 | 32 | 32 |
| | RS19650 | 0.125 | 1 | 32 | 32 |
| | RS19695 | 0.125 | 1 | 32 | 32 |
| | RS20035 | 0.125 | 1 | 32 | 32 |
| | RS20755 | 0.125 | 1 | 32 | 32 |
| | RS21965 | 0.125 | 1 | 32 | 32 |
| | RS25090 | 0.125 | 1 | 32 | 32 |
| | *btuB* (RS25215) | 0.125 | 1 | 32 | 32 |
| | RS27300 | 0.125 | 1 | 32 | 32 |
| | *fptA* (RS28410) | 0.125 | 1 | 32 | 32 |
| | RS29155 | 0.125 | 1 | 32 | 32 |
| | RS29535 | 0.125 | 1 | 32 | **16** |
| | *hmuR* (RS30495) | **0.5** | 1 | 32 | 32 |
| | *orbA2* (RS33275) | 0.125 | 1 | 32 | 32 |
| | RS33410 | 0.125 | 1 | **16** | **16** |
| Regulator | *fur* (RS16355) | **0.03125** | **0.125** | **8** | **4** |
| Energy transduction to TBDRs | *tonB1* (RS20010)** | **0.0625** | **0.25** | 32 | **8** |
| | *tonB2* (RS09735) | 0.125 | 1 | 32 | 32 |
| | *tonB3* (RS07120) | **1** | **2** | 32 | 32 |
| | *tonB4* (RS14515) | 0.125 | 1 | 32 | **16** |

* K562_RS10150 was assigned as a strong homologue to *P. aeruginosa* PAO1 *cntO* but failed reciprocal best-hit BLAST.

** *tonB* paralogues are assigned based on similarity to the *P. aeruginosa* PAO1 *tonB1*

**Table S7.** Bacterial strains and mutants used in this work.

| Strain | Features | Source |
|---|---|---|
| *Achromobacter xylosoxidans* ACH02 | Canadian CF isolate; multidrug resistant | H. Adam (Shared Health MB) |
| *A. xylosoxidans* ACH03 | Canadian CF isolate; multidrug resistant | H. Adam (Shared Health MB) |
| *A. xylosoxidans* ACH09 | Canadian CF isolate; multidrug resistant | H. Adam (Shared Health MB) |
| *B. cenocepacia* 110041 | Canadian CF isolate | National Microbiology Lab, MB |
| *B. cenocepacia* K56-2 | ET12 lineage CF clinical isolate from Toronto | [1] |
| *B. cenocepacia* K56-2::dCas9 | Derived from K56-2; pAH-CTX1rhadCas9 integrated at *attB* site; clean deletion of plasmid accessory genes | [2] |
| *B. cenocepacia* BC7 | Canadian CF isolate; epidemic lineage | CBCCRRR |
| *B. cenocepacia* VC10414 | Canadian CF isolate | CBCCRRR |
| *B. cenocepacia* VC14488 | Canadian CF isolate | CBCCRRR |
| *B. cenocepacia* VC14543 | Canadian CF isolate | CBCCRRR |
| *B. cenocepacia* VC14761 | Canadian CF isolate | CBCCRRR |
| *B. cenocepacia* 110041 | Canadian CF isolate | National Microbiology Lab, Winnipeg, Canada |
| *B. cenocepacia* F4200989 | CF isolate from SickKids Hospital, Toronto, Canada | Dr. Valerie Waters |
| *B. cenocepacia* G1181568-1 | CF isolate from SickKids Hospital, Toronto, Canada | Dr. Valerie Waters |
| *B. cenocepacia* F3071909 | CF isolate from SickKids Hospital, Toronto, Canada | Dr. Valerie Waters |
| *B. cenocepacia* F9141525 | CF isolate from SickKids Hospital, Toronto, Canada | Dr. Valerie Waters |
| *B. cenocepacia* 130723 | Respiratory isolate from Edmonton, Canada | Canadian Antimicrobial Resistance Alliance |
| *B. cenocepacia* 134991 | Respiratory isolate from Halifax, Canada | Canadian Antimicrobial Resistance Alliance |
| *B. cepacia* ATCC 25416 | Onion pathogen, reference strain | ATCC |
| *B. contaminans* 21NG595759 | Canadian CF isolate | H. Adam (Shared Health MB) |
| *B. contaminans* 21NG609698 | Canadian CF isolate | H. Adam (Shared Health MB) |
| *B. contaminans* FFH2055 | Cystic fibrosis clinical isolate from Argentina | [3] |
| *B. contaminans* LMG23361 | Veterinary isolate | BCCM |
| *B. dolosa* CEP021 | Cystic fibrosis isolate | Dr. David Speert |

| | | |
|---|---|---|
| *B. dolosa* AU0645 | CF clinical isolate | Dr. Sarah Kennedy/Dr. Valerie Waters |
| *B. dolosa* LM018941 | CF clinical isolate | Dr. Sarah Kennedy/Dr. Valerie Waters |
| *B. dolosa* AU2167 | CF clinical isolate | Dr. Sarah Kennedy/Dr. Valerie Waters |
| *B. dolosa* AU6940 | CF clinical isolate | Dr. Sarah Kennedy/Dr. Valerie Waters |
| *B. gladioli* VC14812 | Canadian CF isolate | CBCCRRR |
| *B. gladioli* VC19233 | Canadian CF isolate | CBCCRRR |
| *B. gladioli* 113611 | Respiratory isolate from Saskatoon, Canada | Canadian Antimicrobial Resistance Alliance |
| *B. gladioli* 132208 | Respiratory isolate from Montréal, Canada | Canadian Antimicrobial Resistance Alliance |
| *B. multivorans* 130034 | Canadian CF isolate | National Microbiology Lab, MB |
| *B. multivorans* 21NH942533 | Canadian CF isolate | H. Adam (Shared Health MB) |
| *B. multivorans* 21NJ503211 | Canadian CF isolate | H. Adam (Shared Health MB) |
| *B. mutlivorans* ATCC 17616 | Soil isolate | ATCC |
| *B. multivorans* VC15535 | Canadian CF isolate | CBCCRRR |
| *B. multivorans* VC15555 | Canadian CF isolate | CBCCRRR |
| *B. multivorans* VC19694 | Canadian CF isolate | CBCCRRR |
| *B. multivorans* VC9825 | Canadian CF isolate | CBCCRRR |
| *B. multivorans* 131077 | Respiratory isolate from Toronto, Canada | Canadian Antimicrobial Resistance Alliance |
| *B. multivorans* 134955 | Respiratory isolate from Edmonton, Canada | Canadian Antimicrobial Resistance Alliance |
| *B. multivorans* F9091687 | CF clinical isolate | Dr. Sarah Kennedy/Dr. Valerie Waters |
| *B. multivorans* LM013010 | CF clinical isolate | Dr. Sarah Kennedy/Dr. Valerie Waters |

| | | |
|---|---|---|
| *B. multivorans* V1120157 | CF clinical isolate | Dr. Sarah Kennedy/Dr. Valerie Waters |
| *B. multivorans* C5393 | CF clinical isolate | Dr. David Speert |
| *B. thailandensis* E264 | Rice-field soil sample in Thailand | DSMZ |
| *B. vietnamiensis* CEP40 | American cystic fibrosis isolate | [4] |
| *B. vietnamiensis* VC11431 | Canadian cystic fibrosis isolate | CBCCRRR |
| *B. vietnamiensis* VC18984 | Canadian cystic fibrosis isolate | CBCCRRR |
| *E. coli* DH5α | $F^-$ Φ80*lac*ZΔM15 Δ(*lac*ZYA*arg*F) U169 *rec*A1 *end*A1 *hsd*R17 ($rK^-$, $mK^+$) *pho*A *sup*E44 $\lambda^-$ *thi*-1 *gyr*A96 *rel*A1 | Invitrogen |
| *E. coli* MM294 | F− *φ*80 *lac*ZΔM15 *endA1 recA1 hsdR17* (rH− mK+) *supE44 thi-1* Δ*gyrA96* (*ΔlacZYA-argF*)U169 *relA1* | [5] |
| *E. coli* SY327 | $F^-$ *araD* Δ(*lac-proAB*) *argE*(Am) *recA56* Rifr *nalA* λ*pir* | [6] |
| *Pseudomonas. aeruginosa* 20NB681421 | Canadian cystic fibrosis isolate | H. Adam (Shared Health MB) |
| *P. aeruginosa* 21NE72623 | Canadian cystic fibrosis isolate | H. Adam (Shared Health MB) |
| *P. aeruginosa* PAO1 | Lab strain from burn wound | Dr. Ayush Kumar |
| *P. aeruginosa* 70 | CF clinical isolate from Montréal, Canada | Dr. Valerie Waters |
| *P. aeruginosa* 71 | CF clinical isolate from Montréal, Canada | Dr. Valerie Waters |
| *P. aeruginosa* 74 | CF clinical isolate from Montréal, Canada | Dr. Valerie Waters |
| *Stenotrophomonas maltophilia* 20NC580771 | Canadian CF isolate | H. Adam (Shared Health MB) |
| *S. maltophilia* 21NE928511 | Canadian CF isolate | H. Adam (Shared Health MB) |
| *S. maltophilia* D457 | Non-CF respiratory isolate from Spain | ATCC |
| *S. maltophilia* K279a | Bloodstream isolate from UK; multidrug resistant | ATCC |
| *S. maltophilia* 03 T20510 | CF isolate from SickKids Hospital, Toronto, Canada | Dr. Valerie Waters |
| *S. maltophilia* 04 T22674 | CF isolate from SickKids Hospital, Toronto, Canada | Dr. Valerie Waters |
| *S. maltophilia* 05 41846 | CF isolate from SickKids Hospital, Toronto, Canada | Dr. Valerie Waters |

**Table S8.** Plasmids used in this work.

| Plasmid | Features | Source |
|---|---|---|
| pDAI-SceI-*sacB* | pDAI-SceI expressing *sacB* | [7,8] |
| pGPI-SceI | $ori_{R6K}$ $Tmp^r$ $mob^+$ carries I-SceI cut site | [9] |
| pSCrhaBoutgfp | pTnMod-OTp′, *rhaR rhaS* $P_{rhaB}$ *e-gfp* | [10] |
| pRBrhaBout | pSCrhaboutgfp derivative<br>$ori_{R6K}$ *dhfr rhaR rhaS* P*rhaB* (also called pRB-rham) | [11] |
| pKS1 | pRBrhaBout with *Nco*I and *Bam*HI restriction sites inserted after *dhfr* | This study |
| pRBrha-Barcode | Pool of pKS1 with the random barcode ligated in the *Nco*I and *Bam*HI sites | This study |
| pRK2013 | $ori_{colE1}$ RK2 derivative $Kan^r$ $mob^+$ $tra^+$ | [12] |
| pSCrhaB2-plus | pSCrhaB2 with $rhaI_1$-$rhaI_1$ permutation of *rhaS* binding sites upstream of $P_{rhaBAD}$ and bacteriophage T7 *gene 10* stem loop inserted upstream of native *rhaBAD* 5′ UTR | [13] |
| pSCB2-sgRNAv2 | Template for pgRNA created by inverse PCR; derived from pSCrhaB2-sgRNA by inverse<br>PCR to remove *rhaS*, *rhaR*, and<br>$P_{rhaB}$ | [2] |
| pgRNA-non-target | Derived from pSCB2-sgRNA; random 20 nt sequence added as gRNA binding region by inverse PCR | [2] |
| p*hldD*+ | pSCrhaB2plus with K56-2 *hldD* gene (K562_RS14060) cloned between *Nde*I and *Xba*I sites | This study |
| p*waaL*+ | pSCrhaB2plus with K56-2 *waaL* gene (K562_RS06530) cloned between *Nde*I and *Xba*I sites | This study |
| p*wbiI*+ | pSCrhaB2plus with K56-2 *wbiI* gene (K562_RS15035) cloned between *Nde*I and *Xba*I sites | This study |
| p*wzm-wzt*+ | pSCrhaB2plus with K56-2 *wzm* and *wzt* genes (K562_RS15080 and K562_RS15085) cloned between *Nde*I and *Xba*I sites | This study |
| pGPI-SceI-*hldD* | pGPI-SceI with a fusion of 360 bp regions immediately upstream and downstream of *hldD* cloned into *Xba*I and *Xma*I sites. Enables deletion of the 3' 796/993 bp of *hldD*. | This study |
| pGPI-SceI-*waaL* | pGPI-SceI with a fusion of 360 bp regions immediately upstream and downstream of *waaL* cloned into *Xba*I and *Xma*I sites. Enables deletion of the 5' 785/1230 bp of *waaL*. | This study |
| pgRNA-32470-1 | pSCB2-sgRNAv2 expressing sgRNA targeting K562_RS32470 | This study |

| | | |
|---|---|---|
| pgRNA-32470-2 | pSCB2-sgRNAv2 expressing sgRNA targeting K562_RS32470 | This study |
| pgRNA-*ampC*1 | pSCB2-sgRNAv2 expressing sgRNA targeting *ampC* | This study |
| pgRNA-*ampC*2 | pSCB2-sgRNAv2 expressing sgRNA targeting *ampC* | This study |
| pgRNA-*ampD*1 | pSCB2-sgRNAv2 expressing sgRNA targeting *ampD* | This study |
| pgRNA-*ampD*2 | pSCB2-sgRNAv2 expressing sgRNA targeting *ampD* | This study |
| pgRNA-*ampG*1 | pSCB2-sgRNAv2 expressing sgRNA targeting *ampG* | This study |
| pgRNA-*ampG*2 | pSCB2-sgRNAv2 expressing sgRNA targeting *ampG* | This study |
| pgRNA-*bacA*1 | pSCB2-sgRNAv2 expressing sgRNA targeting *bacA* | This study |
| pgRNA-*bacA*2 | pSCB2-sgRNAv2 expressing sgRNA targeting *bacA* | This study |
| pgRNA-*hldD*1 | pSCB2-sgRNAv2 expressing sgRNA targeting *hldD* | This study |
| pgRNA-*hldD*2 | pSCB2-sgRNAv2 expressing sgRNA targeting *hldD* | This study |
| pgRNA-*ispDF*1 | pSCB2-sgRNAv2 expressing sgRNA targeting *ispDF* operon | This study |
| pgRNA-*ispDF*2 | pSCB2-sgRNAv2 expressing sgRNA targeting *ispDF* operon | This study |
| pgRNA-*mlaFEDvacJ*1 | pSCB2-sgRNAv2 expressing sgRNA targeting the *mlaFEDvacJ* operon | This study |
| pgRNA-*mlaFEDvacJ*2 | pSCB2-sgRNAv2 expressing sgRNA targeting the *mlaFEDvacJ* operon | This study |
| pgRNA-*mlaCB*1 | pSCB2-sgRNAv2 expressing sgRNA targeting the *mlaCB* operon | This study |
| pgRNA-*mlaCB*2 | pSCB2-sgRNAv2 expressing sgRNA targeting the *mlaCB* operon | This study |
| pgRNA-*ogcABEI*1 | pSCB2-sgRNAv2 expressing sgRNA targeting *ogcABEI* operon | This study |
| pgRNA-*ogcABEI*2 | pSCB2-sgRNAv2 expressing sgRNA targeting *ogcABEI* operon | This study |
| pgRNA-*penB*1 | pSCB2-sgRNAv2 expressing sgRNA targeting *penB* | This study |
| pgRNA-*penB*2 | pSCB2-sgRNAv2 expressing sgRNA targeting *penB* | This study |
| pgRNA-*penR*1 | pSCB2-sgRNAv2 expressing sgRNA targeting *penR* | This study |
| pgRNA-*penR*2 | pSCB2-sgRNAv2 expressing sgRNA targeting *penR* | This study |
| pgRNA-*pglL*1 | pSCB2-sgRNAv2 expressing sgRNA targeting *pglL* | This study |
| pgRNA-*pglL*2 | pSCB2-sgRNAv2 expressing sgRNA targeting *pglL* | This study |
| pgRNA-*piuA*1 | pSCB2-sgRNAv2 expressing sgRNA targeting *piuA* | This study |
| pgRNA-*piuA*2 | pSCB2-sgRNAv2 expressing sgRNA targeting *piuA* | This study |
| pgRNA-*uppS*1 | pSCB2-sgRNAv2 expressing sgRNA targeting *uppS* | This study |
| pgRNA-*uppS*2 | pSCB2-sgRNAv2 expressing sgRNA targeting *uppS* | This study |
| pgRNA-*vio-wbx*1 | pSCB2-sgRNAv2 expressing sgRNA targeting *vio-wbx* operon | This study |

| | | |
|---|---|---|
| pgRNA-*vio-wbx*2 | pSCB2-sgRNAv2 expressing sgRNA targeting *vio-wbx* operon | This study |
| pgRNA-*waaL*1 | pSCB2-sgRNAv2 expressing sgRNA targeting *waaL* | This study |
| pgRNA-*waaL*2 | pSCB2-sgRNAv2 expressing sgRNA targeting *waaL* | This study |
| pgRNA-*wabR-P*1 | pSCB2-sgRNAv2 expressing sgRNA targeting *wabR-P* operon | This study |
| pgRNA-*wabR-P*2 | pSCB2-sgRNAv2 expressing sgRNA targeting *wabR-P* operon | This study |
| pgRNA-*wbiFGHI*1 | pSCB2-sgRNAv2 expressing sgRNA targeting *wbiFGHI* operon | This study |
| pgRNA-*wbiFGHI*2 | pSCB2-sgRNAv2 expressing sgRNA targeting *wbiFGHI* operon | This study |
| pgRNA-*wzm-wzt*1 | pSCB2-sgRNAv2 expressing sgRNA targeting *wzm-wzt* operon | This study |
| pgRNA-*wzm-wzt*2 | pSCB2-sgRNAv2 expressing sgRNA targeting *wzm-wzt* operon | This study |
| pgRNA-00600-1 | pSCB2-sgRNAv2 expressing sgRNA targeting K562_RS00600 | This study |
| pgRNA-00600-2 | pSCB2-sgRNAv2 expressing sgRNA targeting K562_RS00600 | This study |
| pgRNA-07175-1 | pSCB2-sgRNAv2 expressing sgRNA targeting K562_RS07175 | This study |
| pgRNA-07175-2 | pSCB2-sgRNAv2 expressing sgRNA targeting K562_RS07175 | This study |
| pgRNA-09770-1 | pSCB2-sgRNAv2 expressing sgRNA targeting K562_RS09770 | This study |
| pgRNA-09770-2 | pSCB2-sgRNAv2 expressing sgRNA targeting K562_RS09770 | This study |
| pgRNA-09805-1 | pSCB2-sgRNAv2 expressing sgRNA targeting K562_RS09805 | This study |
| pgRNA-09805-2 | pSCB2-sgRNAv2 expressing sgRNA targeting K562_RS09805 | This study |
| pgRNA-10150-1 | pSCB2-sgRNAv2 expressing sgRNA targeting K562_RS10150 | This study |
| pgRNA-10150-2 | pSCB2-sgRNAv2 expressing sgRNA targeting K562_RS0150 | This study |
| pgRNA-10200-1 | pSCB2-sgRNAv2 expressing sgRNA targeting K562_RS10200 | This study |
| pgRNA-10200-2 | pSCB2-sgRNAv2 expressing sgRNA targeting K562_RS10200 | This study |

| | | |
|---|---|---|
| pgRNA-11910-1 | pSCB2-sgRNAv2 expressing sgRNA targeting K562_RS11910 | This study |
| pgRNA-11910-2 | pSCB2-sgRNAv2 expressing sgRNA targeting K562_RS11910 | This study |
| pgRNA-12040-1 | pSCB2-sgRNAv2 expressing sgRNA targeting K562_RS12040 | This study |
| pgRNA-12040-2 | pSCB2-sgRNAv2 expressing sgRNA targeting K562_RS12040 | This study |
| pgRNA-14405-1 | pSCB2-sgRNAv2 expressing sgRNA targeting K562_RS14405 | This study |
| pgRNA-14405-2 | pSCB2-sgRNAv2 expressing sgRNA targeting K562_RS14405 | This study |
| pgRNA-19650-1 | pSCB2-sgRNAv2 expressing sgRNA targeting K562_RS19650 | This study |
| pgRNA-19650-2 | pSCB2-sgRNAv2 expressing sgRNA targeting K562_RS19650 | This study |
| pgRNA-19695-1 | pSCB2-sgRNAv2 expressing sgRNA targeting K562_RS19695 | This study |
| pgRNA-19695-2 | pSCB2-sgRNAv2 expressing sgRNA targeting K562_RS19695 | This study |
| pgRNA-20035-1 | pSCB2-sgRNAv2 expressing sgRNA targeting K562_RS20035 | This study |
| pgRNA-20035-2 | pSCB2-sgRNAv2 expressing sgRNA targeting K562_RS20035 | This study |
| pgRNA-20755-1 | pSCB2-sgRNAv2 expressing sgRNA targeting K562_RS20755 | This study |
| pgRNA-20755-2 | pSCB2-sgRNAv2 expressing sgRNA targeting K562_RS20755 | This study |
| pgRNA-21965-1 | pSCB2-sgRNAv2 expressing sgRNA targeting K562_RS21965 | This study |
| pgRNA-21965-2 | pSCB2-sgRNAv2 expressing sgRNA targeting K562_RS21965 | This study |
| pgRNA-25090-1 | pSCB2-sgRNAv2 expressing sgRNA targeting K562_RS25090 | This study |
| pgRNA-25090-2 | pSCB2-sgRNAv2 expressing sgRNA targeting K562_RS25090 | This study |
| pgRNA-25215-1 | pSCB2-sgRNAv2 expressing sgRNA targeting K562_RS25215 | This study |
| pgRNA-25215-2 | pSCB2-sgRNAv2 expressing sgRNA targeting K562_RS25215 | This study |

| | | |
|---|---|---|
| pgRNA-27300-1 | pSCB2-sgRNAv2 expressing sgRNA targeting K562_RS27300 | This study |
| pgRNA-27300-2 | pSCB2-sgRNAv2 expressing sgRNA targeting K562_RS27300 | This study |
| pgRNA-28410-1 | pSCB2-sgRNAv2 expressing sgRNA targeting K562_RS28410 | This study |
| pgRNA-28410-2 | pSCB2-sgRNAv2 expressing sgRNA targeting K562_RS28410 | This study |
| pgRNA-29155-1 | pSCB2-sgRNAv2 expressing sgRNA targeting K562_RS29155 | This study |
| pgRNA-29155-2 | pSCB2-sgRNAv2 expressing sgRNA targeting K562_RS29155 | This study |
| pgRNA-29535-1 | pSCB2-sgRNAv2 expressing sgRNA targeting K562_RS29535 | This study |
| pgRNA-29535-2 | pSCB2-sgRNAv2 expressing sgRNA targeting K562_RS29535 | This study |
| pgRNA-30495-1 | pSCB2-sgRNAv2 expressing sgRNA targeting K562_RS30495 | This study |
| pgRNA-30495-2 | pSCB2-sgRNAv2 expressing sgRNA targeting K562_RS30495 | This study |
| pgRNA-33275-1 | pSCB2-sgRNAv2 expressing sgRNA targeting K562_RS33275 | This study |
| pgRNA-33275-2 | pSCB2-sgRNAv2 expressing sgRNA targeting K562_RS33275 | This study |
| pgRNA-33410-1 | pSCB2-sgRNAv2 expressing sgRNA targeting K562_RS33410 | This study |
| pgRNA-33410-2 | pSCB2-sgRNAv2 expressing sgRNA targeting K562_RS33410 | This study |
| pgRNA-16355-1 | pSCB2-sgRNAv2 expressing sgRNA targeting K562_RS16355 | This study |
| pgRNA-16355-2 | pSCB2-sgRNAv2 expressing sgRNA targeting K562_RS16355 | This study |
| pgRNA-20010-1 | pSCB2-sgRNAv2 expressing sgRNA targeting K562_RS20010 | This study |
| pgRNA-20010-2 | pSCB2-sgRNAv2 expressing sgRNA targeting K562_RS20010 | This study |
| pgRNA-09735-1 | pSCB2-sgRNAv2 expressing sgRNA targeting K562_RS09735 | This study |
| pgRNA-09735-2 | pSCB2-sgRNAv2 expressing sgRNA targeting K562_RS09735 | This study |

| | | |
|---|---|---|
| pgRNA-07120-1 | pSCB2-sgRNAv2 expressing sgRNA targeting K562_RS07120 | This study |
| pgRNA-07120-2 | pSCB2-sgRNAv2 expressing sgRNA targeting K562_RS07120 | This study |
| pgRNA-14515-1 | pSCB2-sgRNAv2 expressing sgRNA targeting K562_RS14515 | This study |
| pgRNA-14515-2 | pSCB2-sgRNAv2 expressing sgRNA targeting K562_RS14515 | This study |

**Table S9.** Primers used in this work.

| Primer | Sequence (restriction sites in lowercase) | Notes |
|---|---|---|
| 681 | CAAGCAGAAGACGGCATACGAGATTCGCCTTAGTCTCGTG G GCTCGGAGATGTGTATAAGAGACAG | Used for TnSeq circle |
| 682 | CACAAGTGCGGCCGCACTAGTCTAGATTTAAATTACCGTA GTGAGTTCTTCGTCCGAGCCAC | Collector probe for TnSeq-circle |
| 683 | P – CCGTAGTGAGTTCTTCGTCCGAGCCACTCGGAGATGTGTA T AAGAGACAGT | Half of adapter for TnSeq-circle |
| 684 | P – CTGTCTCTTATACACATCTCCGAGTGGCTCGGACGAAGAA C TCACTACGG | Half of adapter for TnSeq-circle |
| 690 | AATGATACGGCGACCACCGAGATCTACACTAGATCGCTCG TCGGCAGCGTCAGATGTGTATAAGAGACAGNNNNAATCT AGACTAGTGCGGCC | Used for TnSeq circle |
| 715 | CAAGCAGAAGACGGCATACGAGATCTAGTACGGTCTCGTG G GCTCGGAGATGTGTATAAGAGACAG | Used for TnSeq circle |
| 717 | CAAGCAGAAGACGGCATACGAGATGCTCAGGAGTCTCGT GG GCTCGGAGATGTGTATAAGAGACAG | Used for TnSeq circle |
| 718 | CAAGCAGAAGACGGCATACGAGATAGGAGTCCGTCTCGT GG GCTCGGAGATGTGTATAAGAGACAG | Used for TnSeq circle |
| 719 | AATGATACGGCGACCACCGAGATCTACACCTCTCTATTCG TCGGCAGCGTCAGATGTGTATAAGAGACAGNNNNAATCT AGACTAGTGCGGCC | Used for TnSeq circle |
| 729 | CAAGCAGAAGACGGCATACGAGATCATGCCTAGTCTCGTG G GCTCGGAGATGTGTATAAGAGACAG | Used for TnSeq circle |
| 781 | TAAGATggatccTCAGTTGGCTTCATCGCTAC | Colony PCR for pSCB2-sgRNA |
| 847 | TTCCTGTCAGTAACGAGAAGG | Colony PCR for plasmids based on pSCrhaB2; used with 1025 to create pSCB2-sgRNA |
| 848 | CCGCCAGGCAAATTCTGTTT | Reverse primer for colony PCR, all gRNAs |
| 964 | CACTTGTGTATAAGAGTCAG | Reverse primer for In-fusion construction of pKS1 |
| 965 | CTTATACACAAGTGCccatggGTCTTCAACGAGGCAggatccGG CCGCACTAGTCTAGA | Forward primer for In-fusion construction of pKS1 |
| 972 | ATAATACCATGGATGTCCACGAGGTCTCT | Used to amplify the barcode during pRBrha-Barcode construction |

| | | |
|---|---|---|
| 973 | AATTAAGGATCCGTCGACCTGCAGCGTACG | Used to amplify the barcode during pRBrha-Barcode construction |
| 1092 | ACTAGTATTATACCTAGGACTGAGCTAGC | Reverse primer, all gRNAs |
| 1093 | GTTTTAGAGCTAGAAATAGCAAGTTAAAATAAGGC | Forward primer, control gRNA |
| 1409 | GGCTTATGTCAACTGGGTTCG | Forward primer for general colony PCR of sgRNA plasmids |
| 1429 | GGATCCGGCCGCACTAGTCTAGATTTAAATTACCGTAGTGAGTTCTTCGTCCGAGCCAC | Collector probe for RB-TnSeq-circle |
| 2163 | CAAGCAGAAGACGGCATACGAGATGCTTACGGACGTCTCGTGGGCTCGGAGATGTGTATAAGAGACAGGTCGACCTGCAGCGTACG | BarSeq primer with i7 index; UDP0089 |
| 2164 | CAAGCAGAAGACGGCATACGAGATCGCTTGAAGTGTCTCGTGGGCTCGGAGATGTGTATAAGAGACAGGTCGACCTGCAGCGTACG | BarSeq primer with i7 index; UDP0090 |
| 2165 | CAAGCAGAAGACGGCATACGAGATCGCCTTCTGAGTCTCGTGGGCTCGGAGATGTGTATAAGAGACAGGTCGACCTGCAGCGTACG | BarSeq primer with i7 index; UDP0091 |
| 2166 | CAAGCAGAAGACGGCATACGAGATCTGGATATGTGTCTCGTGGGCTCGGAGATGTGTATAAGAGACAGGTCGACCTGCAGCGTACG | BarSeq primer with i7 index; UDP0092 |
| 2167 | CAAGCAGAAGACGGCATACGAGATATACCAACGCGTCTCGTGGGCTCGGAGATGTGTATAAGAGACAGGTCGACCTGCAGCGTACG | BarSeq primer with i7 index; UDP0093 |
| 2168 | CAAGCAGAAGACGGCATACGAGATCAATCTATGAGTCTCGTGGGCTCGGAGATGTGTATAAGAGACAGGTCGACCTGCAGCGTACG | BarSeq primer with i7 index; UDP0094 |
| 2169 | CAAGCAGAAGACGGCATACGAGATGGTGGAATACGTCTCGTGGGCTCGGAGATGTGTATAAGAGACAGGTCGACCTGCAGCGTACG | BarSeq primer with i7 index; UDP0095 |
| 2170 | CAAGCAGAAGACGGCATACGAGATTGGACGGAGGGTCTCGTGGGCTCGGAGATGTGTATAAGAGACAGGTCGACCTGCAGCGTACG | BarSeq primer with i7 index; UDP0096 |
| 2171 | AATGATACGGCGACCACCGAGATCTACACCGTGTATCTTTCGTCGGCAGCGTCAGATGTGTATAAGAGACAGGATGTCCACGAGGTCTCT | BarSeq primer with i5 index; UDP0089 |
| 2172 | AATGATACGGCGACCACCGAGATCTACACGAACCATGAATCGTCGGCAGCGTCAGATGTGTATAAGAGACAGGATGTCCACGAGGTCTCT | BarSeq primer with i5 index; UDP0090 |
| 2173 | AATGATACGGCGACCACCGAGATCTACACGGCCATCATATCGTCGGCAGCGTCAGATGTGTATAAGAGACAGGATGTCCACGAGGTCTCT | BarSeq primer with i5 index; UDP0091 |

| | | |
|---|---|---|
| 2174 | AATGATACGGCGACCACCGAGATCTACACACATACTTCCTCGTCGGCAGCGTCAGATGTGTATAAGAGACAGGATGTCCACGAGGTCTCT | BarSeq primer with i5 index; UDP0092 |
| 2175 | AATGATACGGCGACCACCGAGATCTACACTATGTGCAATTCGTCGGCAGCGTCAGATGTGTATAAGAGACAGGATGTCCACGAGGTCTCT | BarSeq primer with i5 index; UDP0093 |
| 2176 | AATGATACGGCGACCACCGAGATCTACACGATTAAGGTGTCGTCGGCAGCGTCAGATGTGTATAAGAGACAGGATGTCCACGAGGTCTCT | BarSeq primer with i5 index; UDP0094 |
| 2177 | AATGATACGGCGACCACCGAGATCTACACATGTAGACAATCGTCGGCAGCGTCAGATGTGTATAAGAGACAGGATGTCCACGAGGTCTCT | BarSeq primer with i5 index; UDP0095 |
| 2178 | AATGATACGGCGACCACCGAGATCTACACCACATCGGTGTCGTCGGCAGCGTCAGATGTGTATAAGAGACAGGATGTCCACGAGGTCTCT | BarSeq primer with i5 index; UDP0096 |
| 2208 | CAAGCAGAAGACGGCATACGAGATAGACACATTAGTCTCGTGGGCTCGGAGATGTGTATAAGAGACAGGTCGACCTGCAGCGTACG | BarSeq primer with i7 index; UDP0065 |
| 2209 | AATGATACGGCGACCACCGAGATCTACACGTAAGGCATATCGTCGGCAGCGTCAGATGTGTATAAGAGACAGGATGTCCACGAGGTCTCT | BarSeq primer with i5 index; UDP0065 |
| 2210 | CAAGCAGAAGACGGCATACGAGATGCGTTGGTATGTCTCGTGGGCTCGGAGATGTGTATAAGAGACAGGTCGACCTGCAGCGTACG | BarSeq primer with i7 index; UDP0066 |
| 2211 | AATGATACGGCGACCACCGAGATCTACACAATTGCTGCGTCGTCGGCAGCGTCAGATGTGTATAAGAGACAGGATGTCCACGAGGTCTCT | BarSeq primer with i5 index; UDP0066 |
| 2212 | CAAGCAGAAGACGGCATACGAGATAGCACATCCTGTCTCGTGGGCTCGGAGATGTGTATAAGAGACAGGTCGACCTGCAGCGTACG | BarSeq primer with i7 index; UDP0067 |
| 2213 | AATGATACGGCGACCACCGAGATCTACACTTACAATTCCTCGTCGGCAGCGTCAGATGTGTATAAGAGACAGGATGTCCACGAGGTCTCT | BarSeq primer with i5 index; UDP0067 |
| 2214 | CAAGCAGAAGACGGCATACGAGATTTGTTCCGTGGTCTCGTGGGCTCGGAGATGTGTATAAGAGACAGGTCGACCTGCAGCGTACG | BarSeq primer with i7 index; UDP0068 |
| 2215 | AATGATACGGCGACCACCGAGATCTACACAACCTAGCACTCGTCGGCAGCGTCAGATGTGTATAAGAGACAGGATGTCCACGAGGTCTCT | BarSeq primer with i5 index; UDP0068 |
| 2216 | CAAGCAGAAGACGGCATACGAGATAAGTACTCCAGTCTCGTGGGCTCGGAGATGTGTATAAGAGACAGGTCGACCTGCAGCGTACG | BarSeq primer with i7 index; UDP0069 |
| 2217 | AATGATACGGCGACCACCGAGATCTACACTCTGTGTGGATCGTCGGCAGCGTCAGATGTGTATAAGAGACAGGATGTCCACGAGGTCTCT | BarSeq primer with i5 index; UDP0069 |

| | | |
|---|---|---|
| 2218 | CAAGCAGAAGACGGCATACGAGATACGTCAATACGTCTCGTGGGCTCGGAGATGTGTATAAGAGACAGGTCGACCTGCAGCGTACG | BarSeq primer with i7 index; UDP0070 |
| 2219 | AATGATACGGCGACCACCGAGATCTACACGGAATTCCAATCGTCGGCAGCGTCAGATGTGTATAAGAGACAGGATGTCCACGAGGTCTCT | BarSeq primer with i5 index; UDP0070 |
| 2220 | CAAGCAGAAGACGGCATACGAGATGGTGTACAAGGTCTCGTGGGCTCGGAGATGTGTATAAGAGACAGGTCGACCTGCAGCGTACG | BarSeq primer with i7 index; UDP0071 |
| 2221 | AATGATACGGCGACCACCGAGATCTACACAAGCGCGCTTTCGTCGGCAGCGTCAGATGTGTATAAGAGACAGGATGTCCACGAGGTCTCT | BarSeq primer with i5 index; UDP0071 |
| 2222 | CAAGCAGAAGACGGCATACGAGATCCACCTGTGTGTCTCGTGGGCTCGGAGATGTGTATAAGAGACAGGTCGACCTGCAGCGTACG | BarSeq primer with i7 index; UDP0072 |
| 2223 | AATGATACGGCGACCACCGAGATCTACACTGAGCGTTGTTCGTCGGCAGCGTCAGATGTGTATAAGAGACAGGATGTCCACGAGGTCTCT | BarSeq primer with i5 index; UDP0072 |
| 2224 | CAAGCAGAAGACGGCATACGAGATGTTCCGCAGGGTCTCGTGGGCTCGGAGATGTGTATAAGAGACAGGTCGACCTGCAGCGTACG | BarSeq primer with i7 index; UDP0073 |
| 2225 | AATGATACGGCGACCACCGAGATCTACACATCATAGGCTTCGTCGGCAGCGTCAGATGTGTATAAGAGACAGGATGTCCACGAGGTCTCT | BarSeq primer with i5 index; UDP0073 |
| 2226 | CAAGCAGAAGACGGCATACGAGATACCTTATGAAGTCTCGTGGGCTCGGAGATGTGTATAAGAGACAGGTCGACCTGCAGCGTACG | BarSeq primer with i7 index; UDP0074 |
| 2227 | AATGATACGGCGACCACCGAGATCTACACTGTTAGAAGGTCGTCGGCAGCGTCAGATGTGTATAAGAGACAGGATGTCCACGAGGTCTCT | BarSeq primer with i5 index; UDP0074 |
| 2228 | CAAGCAGAAGACGGCATACGAGATCGCTGCAGAGGTCTCGTGGGCTCGGAGATGTGTATAAGAGACAGGTCGACCTGCAGCGTACG | BarSeq primer with i7 index; UDP0075 |
| 2229 | AATGATACGGCGACCACCGAGATCTACACGATGGATGTATCGTCGGCAGCGTCAGATGTGTATAAGAGACAGGATGTCCACGAGGTCTCT | BarSeq primer with i5 index; UDP0075 |
| 2230 | CAAGCAGAAGACGGCATACGAGATGTAGAGTCAGGTCTCGTGGGCTCGGAGATGTGTATAAGAGACAGGTCGACCTGCAGCGTACG | BarSeq primer with i7 index; UDP0076 |
| 2231 | AATGATACGGCGACCACCGAGATCTACACACGGCCGTCATCGTCGGCAGCGTCAGATGTGTATAAGAGACAGGATGTCCACGAGGTCTCT | BarSeq primer with i5 index; UDP0076 |
| 2232 | CAAGCAGAAGACGGCATACGAGATGGATACCAGAGTCTCGTGGGCTCGGAGATGTGTATAAGAGACAGGTCGACCTGCAGCGTACG | BarSeq primer with i7 index; UDP0077 |

| | | |
|---|---|---|
| 2233 | AATGATACGGCGACCACCGAGATCTACACCGTTGCTTACTCGTCGGCAGCGTCAGATGTGTATAAGAGACAGGATGTCCACGAGGTCTCT | BarSeq primer with i5 index; UDP0077 |
| 2234 | CAAGCAGAAGACGGCATACGAGATCGCACTAATGGTCTCGTGGGCTCGGAGATGTGTATAAGAGACAGGTCGACCTGCAGCGTACG | BarSeq primer with i7 index; UDP0078 |
| 2235 | AATGATACGGCGACCACCGAGATCTACACTGACTACATATCGTCGGCAGCGTCAGATGTGTATAAGAGACAGGATGTCCACGAGGTCTCT | BarSeq primer with i5 index; UDP0078 |
| 2236 | CAAGCAGAAGACGGCATACGAGATTCCTGACCGTGTCTCGTGGGCTCGGAGATGTGTATAAGAGACAGGTCGACCTGCAGCGTACG | BarSeq primer with i7 index; UDP0079 |
| 2237 | AATGATACGGCGACCACCGAGATCTACACCGGCCTCGTTTCGTCGGCAGCGTCAGATGTGTATAAGAGACAGGATGTCCACGAGGTCTCT | BarSeq primer with i5 index; UDP0079 |
| 2238 | CAAGCAGAAGACGGCATACGAGATCTGGCTTGCCGTCTCGTGGGCTCGGAGATGTGTATAAGAGACAGGTCGACCTGCAGCGTACG | BarSeq primer with i7 index; UDP0080 |
| 2239 | AATGATACGGCGACCACCGAGATCTACACCAAGCATCCGTCGTCGGCAGCGTCAGATGTGTATAAGAGACAGGATGTCCACGAGGTCTCT | BarSeq primer with i5 index; UDP0080 |
| 2240 | CAAGCAGAAGACGGCATACGAGATACCAGCGACAGTCTCGTGGGCTCGGAGATGTGTATAAGAGACAGGTCGACCTGCAGCGTACG | BarSeq primer with i7 index; UDP0081 |
| 2241 | AATGATACGGCGACCACCGAGATCTACACTCGTCTGACTTCGTCGGCAGCGTCAGATGTGTATAAGAGACAGGATGTCCACGAGGTCTCT | BarSeq primer with i5 index; UDP0081 |
| 2242 | CAAGCAGAAGACGGCATACGAGATTTGTAACGGTGTCTCGTGGGCTCGGAGATGTGTATAAGAGACAGGTCGACCTGCAGCGTACG | BarSeq primer with i7 index; UDP0082 |
| 2243 | AATGATACGGCGACCACCGAGATCTACACCTCATAGCGATCGTCGGCAGCGTCAGATGTGTATAAGAGACAGGATGTCCACGAGGTCTCT | BarSeq primer with i5 index; UDP0082 |
| 2244 | CAAGCAGAAGACGGCATACGAGATGTAAGGCATAGTCTCGTGGGCTCGGAGATGTGTATAAGAGACAGGTCGACCTGCAGCGTACG | BarSeq primer with i7 index; UDP0083 |
| 2245 | AATGATACGGCGACCACCGAGATCTACACAGACACATTATCGTCGGCAGCGTCAGATGTGTATAAGAGACAGGATGTCCACGAGGTCTCT | BarSeq primer with i5 index; UDP0083 |
| 2246 | CAAGCAGAAGACGGCATACGAGATGTCCACTTGTGTCTCGTGGGCTCGGAGATGTGTATAAGAGACAGGTCGACCTGCAGCGTACG | BarSeq primer with i7 index; UDP0084 |
| 2247 | AATGATACGGCGACCACCGAGATCTACACGCGCGATGTTTCGTCGGCAGCGTCAGATGTGTATAAGAGACAGGATGTCCACGAGGTCTCT | BarSeq primer with i5 index; UDP0084 |

| | | |
|---|---|---|
| 2248 | CAAGCAGAAGACGGCATACGAGATTTAGGTACCAGTCTCGTGGGCTCGGAGATGTGTATAAGAGACAGGTCGACCTGCAGCGTACG | BarSeq primer with i7 index; UDP0085 |
| 2249 | AATGATACGGCGACCACCGAGATCTACACCATGAGTACTTCGTCGGCAGCGTCAGATGTGTATAAGAGACAGGATGTCCACGAGGTCTCT | BarSeq primer with i5 index; UDP0085 |
| 2250 | CAAGCAGAAGACGGCATACGAGATGGAATTCCAAGTCTCGTGGGCTCGGAGATGTGTATAAGAGACAGGTCGACCTGCAGCGTACG | BarSeq primer with i7 index; UDP0086 |
| 2251 | AATGATACGGCGACCACCGAGATCTACACACGTCAATACTCGTCGGCAGCGTCAGATGTGTATAAGAGACAGGATGTCCACGAGGTCTCT | BarSeq primer with i5 index; UDP0086 |
| 2252 | CAAGCAGAAGACGGCATACGAGATCATGTAGAGGGTCTCGTGGGCTCGGAGATGTGTATAAGAGACAGGTCGACCTGCAGCGTACG | BarSeq primer with i7 index; UDP0087 |
| 2253 | AATGATACGGCGACCACCGAGATCTACACGATACCTCCTTCGTCGGCAGCGTCAGATGTGTATAAGAGACAGGATGTCCACGAGGTCTCT | BarSeq primer with i5 index; UDP0087 |
| 2254 | CAAGCAGAAGACGGCATACGAGATTACACGCTCCGTCTCGTGGGCTCGGAGATGTGTATAAGAGACAGGTCGACCTGCAGCGTACG | BarSeq primer with i7 index; UDP0088 |
| 2255 | AATGATACGGCGACCACCGAGATCTACACATCCGTAAGTTCGTCGGCAGCGTCAGATGTGTATAAGAGACAGGATGTCCACGAGGTCTCT | BarSeq primer with i5 index; UDP0088 |
| 2421 | GAGCGGGGTCGGGCAAGCAGGTTTTAGAGCTAGAAATAGCAAGTTAAAATAAGGC | Primer containing sgRNA to target *ogcABEI* operon |
| 2422 | TCGTCGTGCCCGTTGTAGGCGTTTTAGAGCTAGAAATAGCAAGTTAAAATAAGGC | Primer containing sgRNA to target *ogcABEI* operon |
| 2445 | TGCGGCGACCAGCAGGGTCGGTTTTAGAGCTAGAAATAGCAAGTTAAAATAAGGC | Primer containing sgRNA to target *ampC* |
| 2446 | TCTGGCGCGTGACGGTGTCGGTTTTAGAGCTAGAAATAGCAAGTTAAAATAAGGC | Primer containing sgRNA to target *ampC* |
| 2453 | GGCCTCGTCTTGAACTGGGCGTTTTAGAGCTAGAAATAGCAAGTTAAAATAAGGC | Primer containing sgRNA to target *piuA* |
| 2454 | GTTTCGGCGGAGGCGAGCAGGTTTTAGAGCTAGAAATAGCAAGTTAAAATAAGGC | Primer containing sgRNA to target *piuA* |
| 2457 | CGTCGCGGCGGCCAGCAACAGTTTTAGAGCTAGAAATAGCAAGTTAAAATAAGGC | Primer containing sgRNA to target *penB* |
| 2458 | GCGGTGGCGGTGAGAACGAGGTTTTAGAGCTAGAAATAGCAAGTTAAAATAAGGC | Primer containing sgRNA to target *penB* |
| 2459 | CGTCACGCTCAGCTCGAGGCGTTTTAGAGCTAGAAATAGCAAGTTAAAATAAGGC | Primer containing sgRNA to target *penR* |
| 2460 | CTGAACGCGTCGCTCAGCACGTTTTAGAGCTAGAAATAGCAAGTTAAAATAAGGC | Primer containing sgRNA to target *penR* |
| 2461 | GGAATCAGGGCGAAAAGTCGGTTTTAGAGCTAGAAATAGCAAGTTAAAATAAGGC | Primer containing sgRNA to target *ispDF* operon |

| | | |
|---|---|---|
| 2462 | CGTGCCGGCACAGGGAATCAGTTTTAGAGCTAGAAATAGC AAGTTAAAATAAGGC | Primer containing sgRNA to target *ispDF* operon |
| 2467 | GCCGACGTCAGGCACGCGAAGTTTTAGAGCTAGAAATAGC AAGTTAAAATAAGGC | Primer containing sgRNA to target *uppS* operon |
| 2468 | ATGATGATCGCGATGTGACGGTTTTAGAGCTAGAAATAGC AAGTTAAAATAAGGC | Primer containing sgRNA to target *uppS* operon |
| 2469 | GATGTTCGCGCCGATAAAACGTTTTAGAGCTAGAAATAGC AAGTTAAAATAAGGC | Primer containing sgRNA to target *hldD* |
| 2470 | ATTGCGAAACTTGTCTGCGCGTTTTAGAGCTAGAAATAGC AAGTTAAAATAAGGC | Primer containing sgRNA to target *hldD* |
| 2477 | TGTTTCGTCGACGGAATGCGGTTTTAGAGCTAGAAATAGC AAGTTAAAATAAGGC | Primer containing sgRNA to target *wbiFGHI* |
| 2478 | GCGGAGACTGGAGAGCGTCCGTTTTAGAGCTAGAAATAGC AAGTTAAAATAAGGC | Primer containing sgRNA to target *wbiFGHI* operon |
| 2479 | ATCGGCTTGTCGAATACGTCGTTTTAGAGCTAGAAATAGC AAGTTAAAATAAGGC | Primer containing sgRNA to target *vio-wbx* operon |
| 2480 | ATTGGCTTGGTCACATAGATGTTTTAGAGCTAGAAATAGC AAGTTAAAATAAGGC | Primer containing sgRNA to target *vio-wbx* operon |
| 2481 | TGTGGATACGTCCGGGGAAGGTTTTAGAGCTAGAAATAGC AAGTTAAAATAAGGC | Primer containing sgRNA to target *wabR-P* operon |
| 2482 | AGCGGCTTGAACAGCAATCTGTTTTAGAGCTAGAAATAGC AAGTTAAAATAAGGC | Primer containing sgRNA to target *wabR-P* operon |
| 2483 | CGCAACGAGCGACAACGAACGTTTTAGAGCTAGAAATAG CAAGTTAAAATAAGGC | Primer containing sgRNA to target *pglL* |
| 2484 | GTGTGATTCGTGATCGCGTAGTTTTAGAGCTAGAAATAGC AAGTTAAAATAAGGC | Primer containing sgRNA to target *pglL* |
| 2485 | AGGTGACCGGTGCTCGACACGTTTTAGAGCTAGAAATAGC AAGTTAAAATAAGGC | Primer containing sgRNA to target *bacA* |
| 2486 | GCCCGCGACGATCAGGTGACGTTTTAGAGCTAGAAATAGC AAGTTAAAATAAGGC | Primer containing sgRNA to target *bacA* |
| 2619 | ATTAcatatgACCCTCATCGTCACCGG | Used to amplify *hldD* (K562_RS14060) |
| 2620 | ATTAtctagaGACCCAGTCTATCGATGCAACG | Used to amplify *hldD* (K562_RS14060) |
| 2627 | ATTAcatatgTTCGCCATCATGTTCGG | Used to amplify *waaL* (K562_RS06530) |
| 2628 | ATTAtctagaTTCACGATCCGCTCCCC | Used to amplify *waaL* (K562_RS06530) |
| 2629 | ATTAcatatgTTGCAATCCAGAGCATCTTGG | Used to amplify *wbiI* (K562_RS15035) |
| 2630 | ATTAtctagaCCTGGAGCCACGTCACCC | Used to amplify *wbiI* (K562_RS15035) |
| 2631 | ATTAcatatgCGGGATAACATTCAAAGTTCC | Used to amplify *wzm-wzt* (K562_RS15085-80) |
| 2632 | ATTAtctagaCGTCGACGCTAAACTCATCATC | Used to amplify *wzm-wzt* (K562_RS15085-80) |

| | | |
|---|---|---|
| 2672 | AGGCGGCGATGCCGCCAGAGGTTTTAGAGCTAGAAATAG CAAGTTAAAATAAGGC | Primer containing sgRNA to target *wzm-wzt* operon |
| 2673 | ATACGGCCAGCAGGAGCAACGTTTTAGAGCTAGAAATAG CAAGTTAAAATAAGGC | Primer containing sgRNA to target *wzm-wzt* operon |
| 2674 | AACGGCCACCGGAACACCGCGTTTTAGAGCTAGAAATAGC AAGTTAAAATAAGGC | Primer containing sgRNA to target *waaL* |
| 2675 | TCGGGACGAACAACGGCCACGTTTTAGAGCTAGAAATAGC AAGTTAAAATAAGGC | Primer containing sgRNA to target *waaL* |
| 2693 | GCTTTGTCGGTTGCGTGA | Colony PCR for *hldD* deletion |
| 2694 | CACTGTTGCTCACCCGTTC | Colony PCR for *hldD* deletion |
| 2695 | CGCCGATCTTTGGCATCTCTA | Colony PCR for *waaL* deletion |
| 2696 | GTCGGTCGGGAAGGCAAC | Colony PCR for *waaL* deletion |
| 2805 | GCAGGGTCTCGGTAGGAGTGGTTTTAGAGCTAGAAATAGC AAGTTAAAATAAGGC | Primer containing sgRNA to target *mlaFEDvacJ* operon |
| 2806 | GCGAAGTTCCAGCAGGGTCTGTTTTAGAGCTAGAAATAGC AAGTTAAAATAAGGC | Primer containing sgRNA to target *mlaFEDvacJ* operon |
| 2807 | TGTTGACGATCGAGATGATGGTTTTAGAGCTAGAAATAGC AAGTTAAAATAAGGC | Primer containing sgRNA to target *mlaCB* operon |
| 2808 | GTGCGGCGGAAATCGGTGTAGTTTTAGAGCTAGAAATAGC AAGTTAAAATAAGGC | Primer containing sgRNA to target *mlaCB* operon |
| 2847 | TCGTGAAATCCCCGACACGCGTTTTAGAGCTAGAAATAGC AAGTTAAAATAAGGC | Primer containing sgRNA to target K562_RS32470 |
| 2848 | GCGCTGCATCGCGGTAGCGTGTTTTAGAGCTAGAAATAGC AAGTTAAAATAAGGC | Primer containing sgRNA to target K562_RS32470 |
| 3018 | CGCGGCGTGGACGGACAGGCGTTTTAGAGCTAGAAATAG CAAGTTAAAATAAGGC | Primer containing sgRNA to targetK562_RS00600 |
| 3019 | GTTGAGGGCGGGCAGTACCGGTTTTAGAGCTAGAAATAGC AAGTTAAAATAAGGC | Primer containing sgRNA to targetK562_RS00600 |
| 3020 | GGATTCATACGTGGTGCAGCGTTTTAGAGCTAGAAATAGC AAGTTAAAATAAGGC | Primer containing sgRNA to targetK562_RS07120 |
| 3021 | ACCGCGACGACGATCACTCGGTTTTAGAGCTAGAAATAGC AAGTTAAAATAAGGC | Primer containing sgRNA to targetK562_RS07120 |
| 3022 | ATCGGCGTGAGACGCAGGACGTTTTAGAGCTAGAAATAGC AAGTTAAAATAAGGC | Primer containing sgRNA to targetK562_RS07175 |
| 3023 | AGCGCAATGCCGAAGGCAATGTTTTAGAGCTAGAAATAGC AAGTTAAAATAAGGC | Primer containing sgRNA to targetK562_RS07175 |
| 3024 | TGCTTCTTGCCGAATTCCCGGTTTTAGAGCTAGAAATAGC AAGTTAAAATAAGGC | Primer containing sgRNA to targetK562_RS09735 |
| 3025 | ATGCCGCCGAAACGGCGCACGTTTTAGAGCTAGAAATAGC AAGTTAAAATAAGGC | Primer containing sgRNA to targetK562_RS09735 |

| | | |
|---|---|---|
| 3026 | GGCTGGCTCTGGGCGAACGCGTTTTAGAGCTAGAAATAGCAAGTTAAAATAAGGC | Primer containing sgRNA to targetK562_RS09770 |
| 3027 | TCCGCAATCGATGTGGCGCCGTTTTAGAGCTAGAAATAGCAAGTTAAAATAAGGC | Primer containing sgRNA to targetK562_RS09770 |
| 3028 | AGGCGTTGTCTGCGCGTAAGGTTTTAGAGCTAGAAATAGCAAGTTAAAATAAGGC | Primer containing sgRNA to targetK562_RS09805 |
| 3029 | GGTCGCGCTGCCGGCTTGCTGTTTTAGAGCTAGAAATAGCAAGTTAAAATAAGGC | Primer containing sgRNA to targetK562_RS09805 |
| 3030 | GGCAATCGAGCACAGCGACAGTTTTAGAGCTAGAAATAGCAAGTTAAAATAAGGC | Primer containing sgRNA to targetK562_RS10150 |
| 3031 | GGCAGGGCAGGAAGCAACGCGTTTTAGAGCTAGAAATAGCAAGTTAAAATAAGGC | Primer containing sgRNA to targetK562_RS10150 |
| 3032 | GTTCATGGCCGGCGCCGTCTGTTTTAGAGCTAGAAATAGCAAGTTAAAATAAGGC | Primer containing sgRNA to targetK562_RS10200 |
| 3033 | CAGCGTGCCGTTTTGCGCGCGTTTTAGAGCTAGAAATAGCAAGTTAAAATAAGGC | Primer containing sgRNA to targetK562_RS10200 |
| 3034 | GGAAGCCATGATCGGAGAGTGTTTTAGAGCTAGAAATAGCAAGTTAAAATAAGGC | Primer containing sgRNA to targetK562_RS11910 |
| 3035 | AGACGGATGGAAGCCATGATGTTTTAGAGCTAGAAATAGCAAGTTAAAATAAGGC | Primer containing sgRNA to targetK562_RS11910 |
| 3036 | GTCGGCCGACGCGGACGATGGTTTTAGAGCTAGAAATAGCAAGTTAAAATAAGGC | Primer containing sgRNA to targetK562_RS12040 |
| 3037 | AGCCGCGAGCCCGTTTCGAGGTTTTAGAGCTAGAAATAGCAAGTTAAAATAAGGC | Primer containing sgRNA to targetK562_RS12040 |
| 3038 | AGCGGCCGGCCATGCGAAAGGTTTTAGAGCTAGAAATAGCAAGTTAAAATAAGGC | Primer containing sgRNA to targetK562_RS14405 |
| 3039 | GTCGGTCGAAGCCGCGTGAGGTTTTAGAGCTAGAAATAGCAAGTTAAAATAAGGC | Primer containing sgRNA to targetK562_RS14405 |
| 3040 | GATCGTGCACGCGGCAAGCAGTTTTAGAGCTAGAAATAGCAAGTTAAAATAAGGC | Primer containing sgRNA to targetK562_RS14515 |
| 3041 | ATTGCGTTCGGCGACACGATGTTTTAGAGCTAGAAATAGCAAGTTAAAATAAGGC | Primer containing sgRNA to targetK562_RS14515 |
| 3042 | CCGATATTCTTGAGATCCGTGTTTTAGAGCTAGAAATAGCAAGTTAAAATAAGGC | Primer containing sgRNA to targetK562_RS16355 |
| 3043 | GAGAATCTTGAGGCGCGGTAGTTTTAGAGCTAGAAATAGCAAGTTAAAATAAGGC | Primer containing sgRNA to targetK562_RS16355 |
| 3044 | TGTGCGTGGGCCTGCCCGGCGTTTTAGAGCTAGAAATAGCAAGTTAAAATAAGGC | Primer containing sgRNA to targetK562_RS19650 |
| 3045 | TTCCTGTGCGTGGGCCTGCCGTTTTAGAGCTAGAAATAGCAAGTTAAAATAAGGC | Primer containing sgRNA to targetK562_RS19650 |
| 3046 | CGCAAACGAAGTGCGGGCGCGTTTTAGAGCTAGAAATAGCAAGTTAAAATAAGGC | Primer containing sgRNA to targetK562_RS19695 |
| 3047 | CGCCTGCGCAAACGAAGTGCGTTTTAGAGCTAGAAATAGCAAGTTAAAATAAGGC | Primer containing sgRNA to targetK562_RS19695 |
| 3048 | GACCGACAGCGGTCGCGCCCGTTTTAGAGCTAGAAATAGCAAGTTAAAATAAGGC | Primer containing sgRNA to targetK562_RS20010 |

| | | |
|---|---|---|
| 3049 | AATTCGACCGTCATCGGCAAGTTTTAGAGCTAGAAATAGC AAGTTAAAATAAGGC | Primer containing sgRNA to targetK562_RS20010 |
| 3050 | CGTCAACAGTCCCCCCAGCGGTTTTAGAGCTAGAAATAGC AAGTTAAAATAAGGC | Primer containing sgRNA to targetK562_RS20035 |
| 3051 | GCGCCGTCCTCTGCCCACGCGTTTTAGAGCTAGAAATAGC AAGTTAAAATAAGGC | Primer containing sgRNA to targetK562_RS20035 |
| 3052 | GGGCGTGTCGCCCTGGGCCGGTTTTAGAGCTAGAAATAGC AAGTTAAAATAAGGC | Primer containing sgRNA to targetK562_RS20755 |
| 3053 | CGGCGCGGGCGTGTCGCCCTGTTTTAGAGCTAGAAATAGC AAGTTAAAATAAGGC | Primer containing sgRNA to targetK562_RS20755 |
| 3054 | ACGGTGAGTTTCAGCATGCGGTTTTAGAGCTAGAAATAGC AAGTTAAAATAAGGC | Primer containing sgRNA to targetK562_RS21965 |
| 3055 | AGCGCGCCGACAGCCAGCGCGTTTTAGAGCTAGAAATAGC AAGTTAAAATAAGGC | Primer containing sgRNA to targetK562_RS21965 |
| 3056 | GCCGGACGCGCCCGGTGCATGTTTTAGAGCTAGAAATAGC AAGTTAAAATAAGGC | Primer containing sgRNA to targetK562_RS25090 |
| 3057 | AGCGGCGCGGATGCGTTGGCGTTTTAGAGCTAGAAATAGC AAGTTAAAATAAGGC | Primer containing sgRNA to targetK562_RS25090 |
| 3058 | GGCGAGCGGGGAAAGCAGCGGTTTTAGAGCTAGAAATAG CAAGTTAAAATAAGGC | Primer containing sgRNA to targetK562_RS25215 |
| 3059 | TTCGGGCAGGTGCCGCGTGTGTTTTAGAGCTAGAAATAGC AAGTTAAAATAAGGC | Primer containing sgRNA to targetK562_RS25215 |
| 3060 | GAGCTTCAGCTCGTCGGGACGTTTTAGAGCTAGAAATAGC AAGTTAAAATAAGGC | Primer containing sgRNA to targetK562_RS27300 |
| 3061 | AGATGGCCTTCGGTGCTGGCGTTTTAGAGCTAGAAATAGC AAGTTAAAATAAGGC | Primer containing sgRNA to targetK562_RS27300 |
| 3064 | CGGACGCCAGCGCCCGAAACGTTTTAGAGCTAGAAATAGC AAGTTAAAATAAGGC | Primer containing sgRNA to targetK562_RS28410 |
| 3065 | GCATCGGCGAACGCCGCCGCGTTTTAGAGCTAGAAATAGC AAGTTAAAATAAGGC | Primer containing sgRNA to targetK562_RS28410 |
| 3066 | CCGCATCGCGCCGTACCACCGTTTTAGAGCTAGAAATAGC AAGTTAAAATAAGGC | Primer containing sgRNA to targetK562_RS29155 |
| 3067 | GAACAGCGCCAACGTGCCAAGTTTTAGAGCTAGAAATAGC AAGTTAAAATAAGGC | Primer containing sgRNA to targetK562_RS29155 |
| 3068 | GGCACGGCAGGGCGTTCGACGTTTTAGAGCTAGAAATAGC AAGTTAAAATAAGGC | Primer containing sgRNA to targetK562_RS29535 |
| 3069 | ACGGACGTGTGGTCCGGGCAGTTTTAGAGCTAGAAATAGC AAGTTAAAATAAGGC | Primer containing sgRNA to targetK562_RS29535 |
| 3070 | CCGAACAGTGCGGCACAGATGTTTTAGAGCTAGAAATAGC AAGTTAAAATAAGGC | Primer containing sgRNA to targetK562_RS30495 |
| 3071 | GCCGAACGCGCCGAACAGTGGTTTTAGAGCTAGAAATAGC AAGTTAAAATAAGGC | Primer containing sgRNA to targetK562_RS30495 |
| 3072 | GGCCACGCACATTGCCCCGCGTTTTAGAGCTAGAAATAGC AAGTTAAAATAAGGC | Primer containing sgRNA to targetK562_RS33275 |
| 3073 | GCCGGCCCTCGCACGTCGATGTTTTAGAGCTAGAAATAGC AAGTTAAAATAAGGC | Primer containing sgRNA to targetK562_RS33275 |

| | | |
|---|---|---|
| 3074 | CGCGTCGGCGCTGCCCATGCGTTTTAGAGCTAGAAATAGC AAGTTAAAATAAGGC | Primer containing sgRNA to targetK562_RS33410 |
| 3075 | GTCGATTCACGCGTACGAAAGTTTTAGAGCTAGAAATAGC AAGTTAAAATAAGGC | Primer containing sgRNA to targetK562_RS33410 |

**Table S10.** gBlock fragments used in this work to delete *hldD* and *waaL* (restriction sites are in lowercase)

| Name | Sequence (restriction sites in lowercase) | Description |
|---|---|---|
| *hldD* frag | ATTAcccgggATCGTCAAGCCTTCCTGCTGAAGCCGCTCGACGGTGTG<br>CCCGCCCATGCCCTTCACACGCTTGGCGAGATCGTCCGCATTCCTG<br>AACGGACCGCGTGCGCCGCGCTCGTCGAGAATCGCCTTCGCGCGTG<br>CCGGGCCGATGCCCTTGATGCCGACGAGCGCATCCTCGTTCGCGGTG<br>TTGACGTCGACGGCCGCCCAGGCCGACGCTACTGCGCCGAGCATGA<br>CCGCTGCGGCAAACCATTCCCTGAACATGTGCTGGATCTCCGAAGAA<br>AACGGCCGGCGGCTGCCGACCGTGAGACCCAGTCTATCGATGCAAC<br>GCGCCGGTTTAAACCTGGCCGGATAGCCAACGCACGTAACGATCTTG<br>TCGAGATAGTCGTCGATCTCGCAATCGACGAGATTGCGAAACTTGTC<br>TGCGCGGGTCAGGTTGTCGACCGCGATGATGCGCGTCTCGCCGCGC<br>TCGTTGAGCGCCTTGACGATGTTCGCGCCGATAAAACCGGCTGCGCC<br>GGTGACGATGAGGGTCATGATCGTCCTGCCTGAAAAGTGCGAGCCC<br>ACTCGGGCGACGCGCTCGTCGCGCCGCCGAATGCGCTCAGTGAAACA<br>GTTCGTCGTAATCCACCGTGGCCGTGCCGAGCTTGCCGACCACGATG<br>CCTGCCGCGCGATTCGCGAGCACCACCGCGTCGACGAGCGGCACCCC<br>GGCGCCGAGCATCGTCGCGACCGTCGCGATCAtctagaTTA | Enables deletion of the 3' 796bp (of total 993 bp) of *hldD* |
| *waaL* frag | ATTATcccgggTGCCGGACGTTACTGGCGCGACGTTCCGTTCGTCGCC<br>GTAACGATCGCGGACATGCCATGCCGCGATCAACCGTATATTTTCC<br>GGCGTCACGTCCAGTGGGTCAGGCGATGCGATCGCGACGCCACGAC<br>ATGCGTCGCGTGGCACGACGGCTCCGCGGGCTGCATGAGCGGGCT<br>GTCGGTGGACGCGGTACGTGGCGCACTGAAACACGTTCTGGTGCAA<br>GTTCGATGACGGCTATTGCGGCGCGTATGGAGCGTCTTCTGTGGATT<br>GGCTGCCCCGTTCTGATGTTCGCCATCATGTTCGGGCACATGGCGGC<br>GTTCGTGAATTGCAGCATGGTGCTGGTCGGGATTGCGCTGATCGAA<br>ACCGGTGCCGGGAATGGCCTACCGCAGCGAGATGCCGCCCGAATTGC<br>TGGCACTCGAGCCGCAGGCGCTGACGCACGCGCACAATCTTTTCCTG<br>AATACGTGGCTACAGACCGGGCGCATCGGCCTGGCAATGCAGGCACT<br>GTTGCTGGCGGCGCTCGTATTCGCGTTCTGGCGACTGCGGAAGGCTG<br>ACCCGTGGATTGCGGCGGGCGGCATCGCGCTCGTGGTCGGGATGGTC<br>ACGAAGAACCTGGTGGACGATTTCATGTGGCAGACGACGATCCTGGC<br>ATTCTGGGCATTTGCCGGGCTGGTGCTCGGGCATGGCGAGCGACGCG<br>CACGCGTGCGTCGCGCGCAACCGGGGAGCGGATCGTGAAtctagaTTAT | Enables deletion of the 5' 785 bp (of total 1230 bp) of *waaL* |

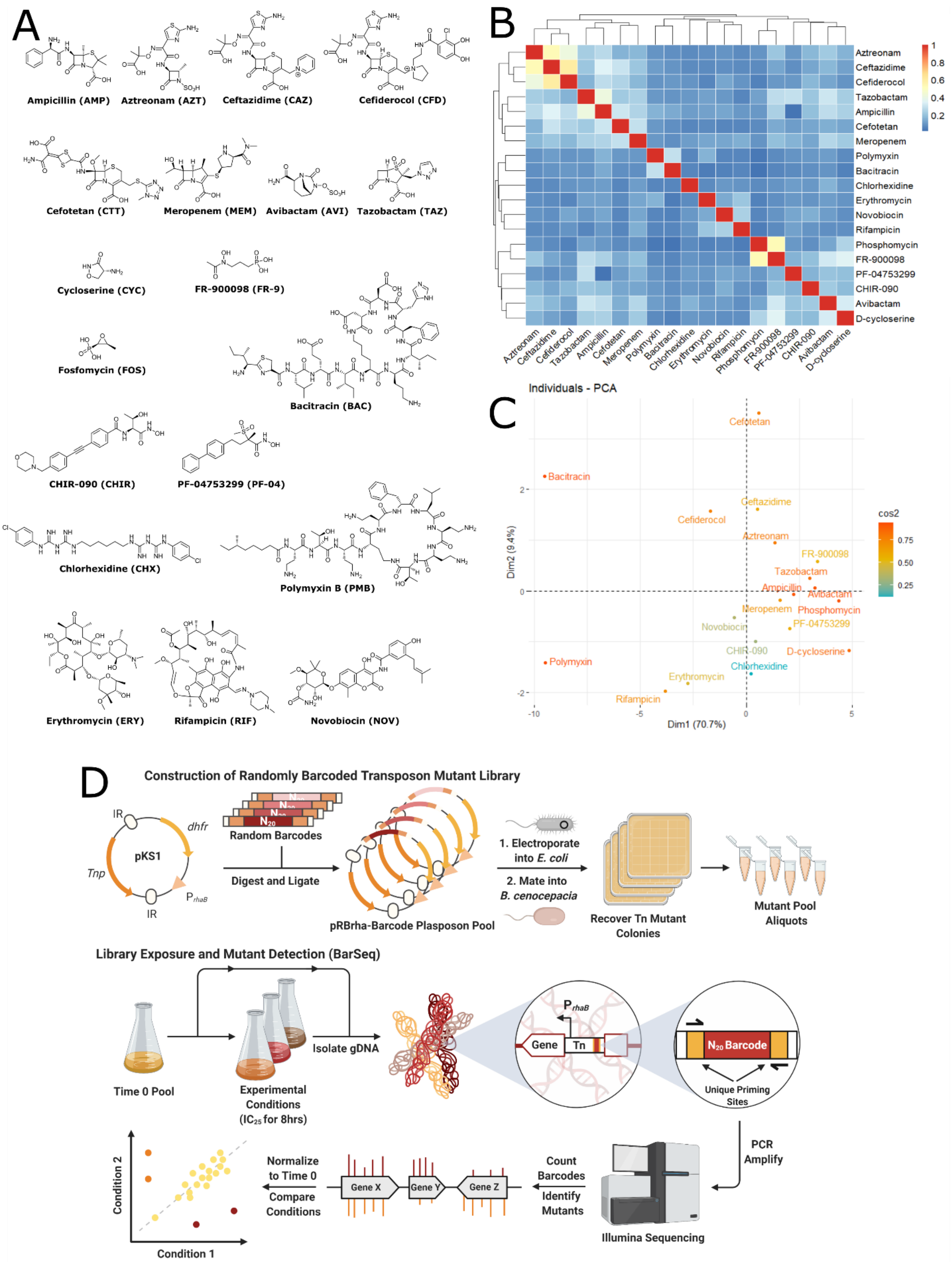

A
Ampicillin (AMP)
Aztreonam (AZT)
Ceftazidime (CAZ)
Cefiderocol (CFD)
Cefotetan (CTT)
Meropenem (MEM)
Avibactam (AVI)
Tazobactam (TAZ)
Cycloserine (CYC)
FR-900098 (FR-9)
Fosfomycin (FOS)
Bacitracin (BAC)
CHIR-090 (CHIR)
PF-04753299 (PF-04)
Chlorhexidine (CHX)
Polymyxin B (PMB)
Erythromycin (ERY)
Rifampicin (RIF)
Novobiocin (NOV)
B
Aztreonam
Ceftazidime
Cefiderocol
Tazobactam
Ampicillin
Cefotetan
Meropenem
Polymyxin
Bacitracin
Chlorhexidine
Erythromycin
Novobiocin
Rifampicin
Phosphomycin
FR-900098
PF-04753299
CHIR-090
Avibactam
D-cycloserine
1
0.8
0.6
0.4
0.2
C
Individuals - PCA
Dim1 (70.7%)
Dim2 (9.4%)
cos2
0.75
0.50
0.25
Cefotetan
Bacitracin
Ceftazidime
Cefiderocol
Aztreonam
FR-900098
Tazobactam
Ampicillin
Avibactam
Meropenem
Phosphomycin
PF-04753299
Novobiocin
CHIR-090
D-cycloserine
Polymyxin
Chlorhexidine
Erythromycin
Rifampicin
D
Construction of Randomly Barcoded Transposon Mutant Library
IR
dhfr
pKS1
Tnp
IR
Random Barcodes
Digest and Ligate
pRBrha-Barcode Plasposon Pool
1. Electroporate into E. coli
2. Mate into B. cenocepacia
Recover Tn Mutant Colonies
Mutant Pool Aliquots
Library Exposure and Mutant Detection (BarSeq)
Time 0 Pool
Experimental Conditions (IC25 for 8hrs)
Isolate gDNA
Gene
Tn
N20 Barcode
Unique Priming Sites
PCR Amplify
Illumina Sequencing
Count Barcodes
Identify Mutants
Gene X
Gene Y
Gene Z
Normalize to Time 0
Compare Conditions
Condition 2
Condition 1

**Figure S1.** Schematic of creating randomly-barcoded transposon mutants and exposure to an antibiotic panel. A) Structure of the antibiotics used in the BarSeq experiment. B) 2-D hierarchical clustering of atom pair Tanimoto similarity scoring from the ChemMine Tools package (Backman et al., 2011). C) Principal component analysis based on chemical properties from the ChemMine Tools package. Warmer colours indicate better representation by the principal components. D). DNA fragments containing random 20 bp barcodes flanked by priming and restriction sites are ligated into the parent plasposon pKS1. The resultant plasposon pool is electroporated into E. coli and then introduced into K56-2 by triparental mating. Mutant colonies are recovered and stored until needed. After growth in a desired condition, genomic DNA is isolated and used as template for one-step PCR to amplify barcodes and add flow cell adapters and sequencing indices. Prepared DNA libraries are then sequenced on an Illumina platform (BarSeq). Barcode abundance is counted and compared between conditions by a bioinformatic pipeline.

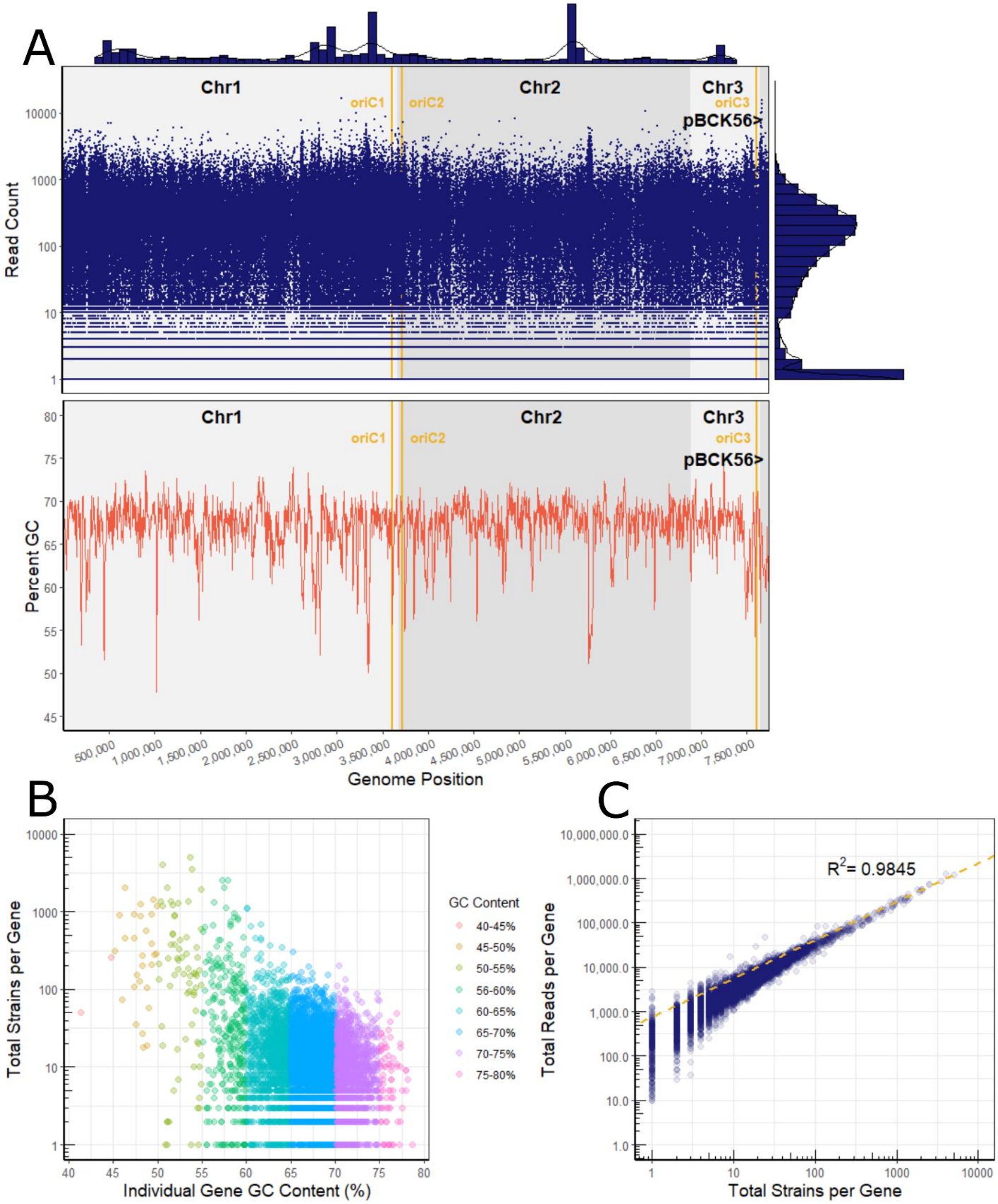
A
Chr1
Chr2
Chr3
oriC1
oriC2
oriC3
pBCK56>
Read Count
Percent GC
Genome Position
B
Total Strains per Gene
Individual Gene GC Content (%)
GC Content
40-45%
45-50%
50-55%
56-60%
60-65%
65-70%
70-75%
75-80%
C
$R^2$= 0.9845
Total Reads per Gene
Total Strains per Gene

**Figure S2**. Transposon insertion density across the K56-2 genome. A). Each insertion in the library was mapped along with the read count. The marginal histograms represent 1D point density in the x and y dimensions of the plot. The x-marginal histogram has a bin width of 100,000 bp and shows insertion density. The y-marginal histogram splits each $log_{10}$ into 5 bins and shows read count density. The chromosomal origins (OriC) were identified by homology from *B. cenocepacia* J2315 using DoriC 10.0 *[14]*. The GC_content perl script (https://github.com/DamienFr/GC_content_in_sliding_window) was used to calculate GC content with a 5,000 bp window and 5,000 bp step size. B) Gene-level breakdown of the number of strains per gene versus GC content. Each point is coloured by binned GC-content of the gene. C) The number of reads per gene versus the number of strains the gene is represented by in the library. The dashed yellow line is a linear regression showing the relationship between the number of reads and number of strains for each gene.

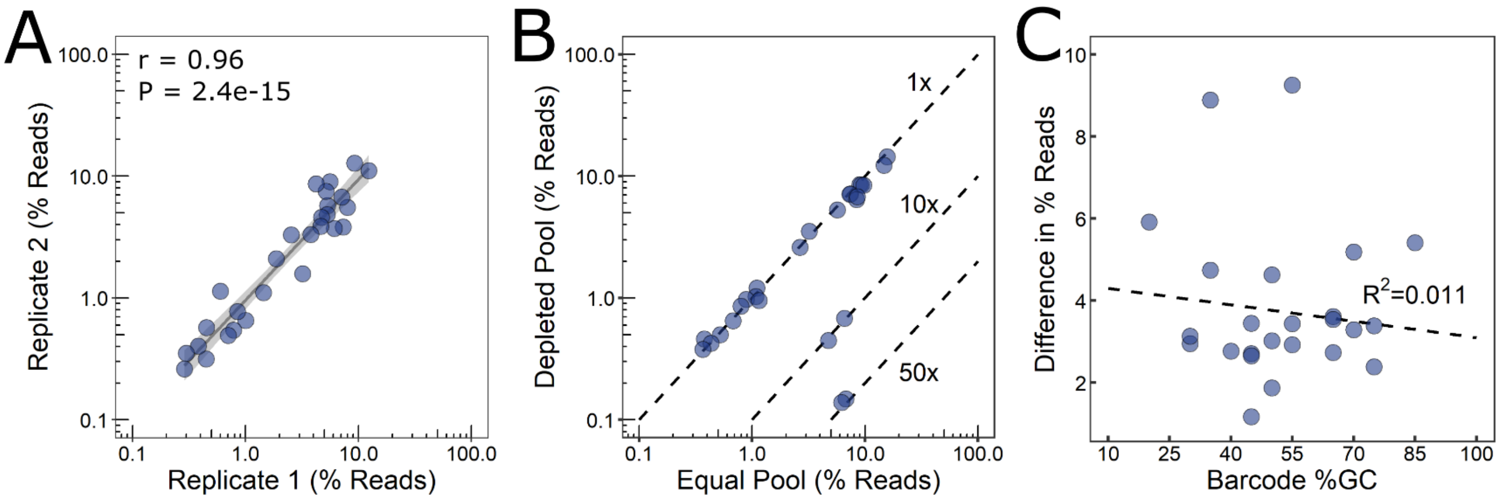


**Figure S3**. Validation of BarSeq to reproducibly and accurately determine barcoded transposon mutant abundance. A) Reproducibility of BarSeq shown between two replicates of artificial equal pools of 30 mutants. The grey line shows a linear regression with 95% confidence interval in lighter grey. The r and P-values are from Pearson's correlation test. B) Detection of changes in barcode abundance in artificial pools of 30 mutants (equal vs depleted at 10x and 50x). Each point is an average of two replicates. Dashed lines indicate expected mutant abundance in the two pools: 1x, 10x, or 50x. C) Absolute difference between the observed and expected barcode abundance (in percent of reads) for the equal pool replicates as a function of individual barcode GC-content. If truly equal, all barcodes would be present at ~3%. The dashed line shows a linear regression.

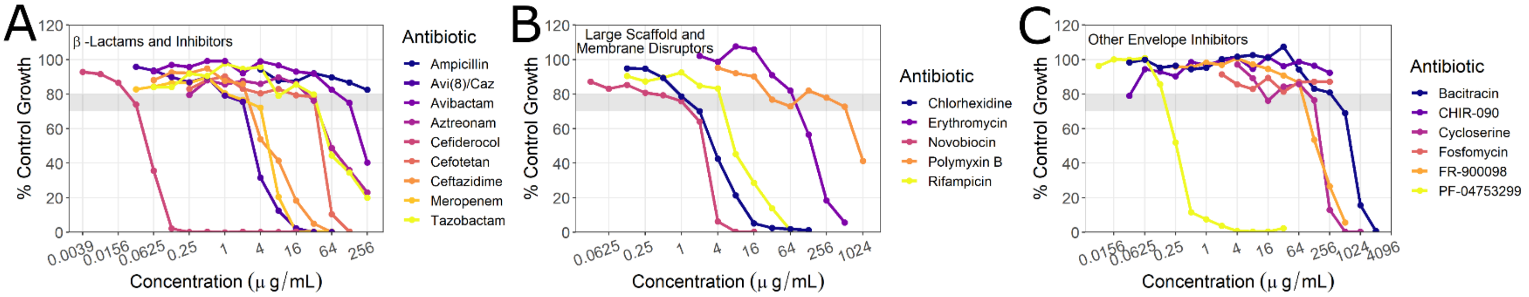


**Figure S4.** Growth dose-response curves of wild-type K56-2 to antibiotics in the panel. The grey box indicates the target region of 20 – 30% growth inhibition relative to the no-antibiotic control. Points are averages of three replicates. Cells were grown in LB to more closely represent conditions of the mutant library exposure.

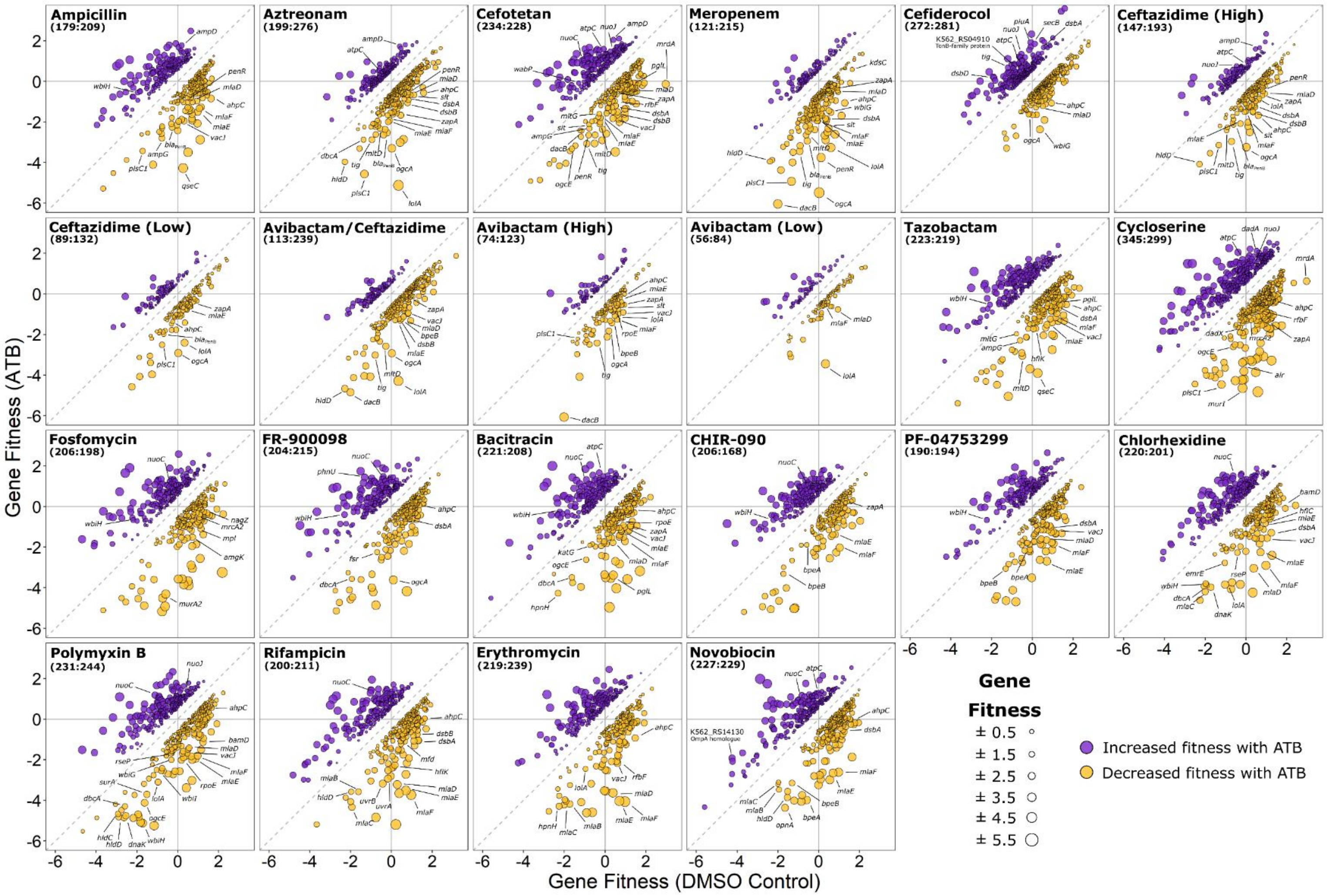


**Figure S5.** Gene fitness profiles of K56-2 transposon mutant library exposed to cell envelope-targeting antibiotic panel. Each point represents a gene that when disrupted by a transposon, affected fitness of the mutant in the presence of an antibiotic versus the DMSO control (two-sided t-test; $P < 0.05$; only genes with fitness effects greater than 0.5 or less than -0.5 were considered). The points are coloured based on positive (increased fitness; purple) or negative (decreased fitness; gold) interactions between the genes and tested antibiotics. The absolute difference in fitness score for a gene in each condition versus the DMSO control is given by the size of the point. Select genes of interest are indicated and named. The total number of genes in each condition with altered fitness (increased:decreased) is shown.

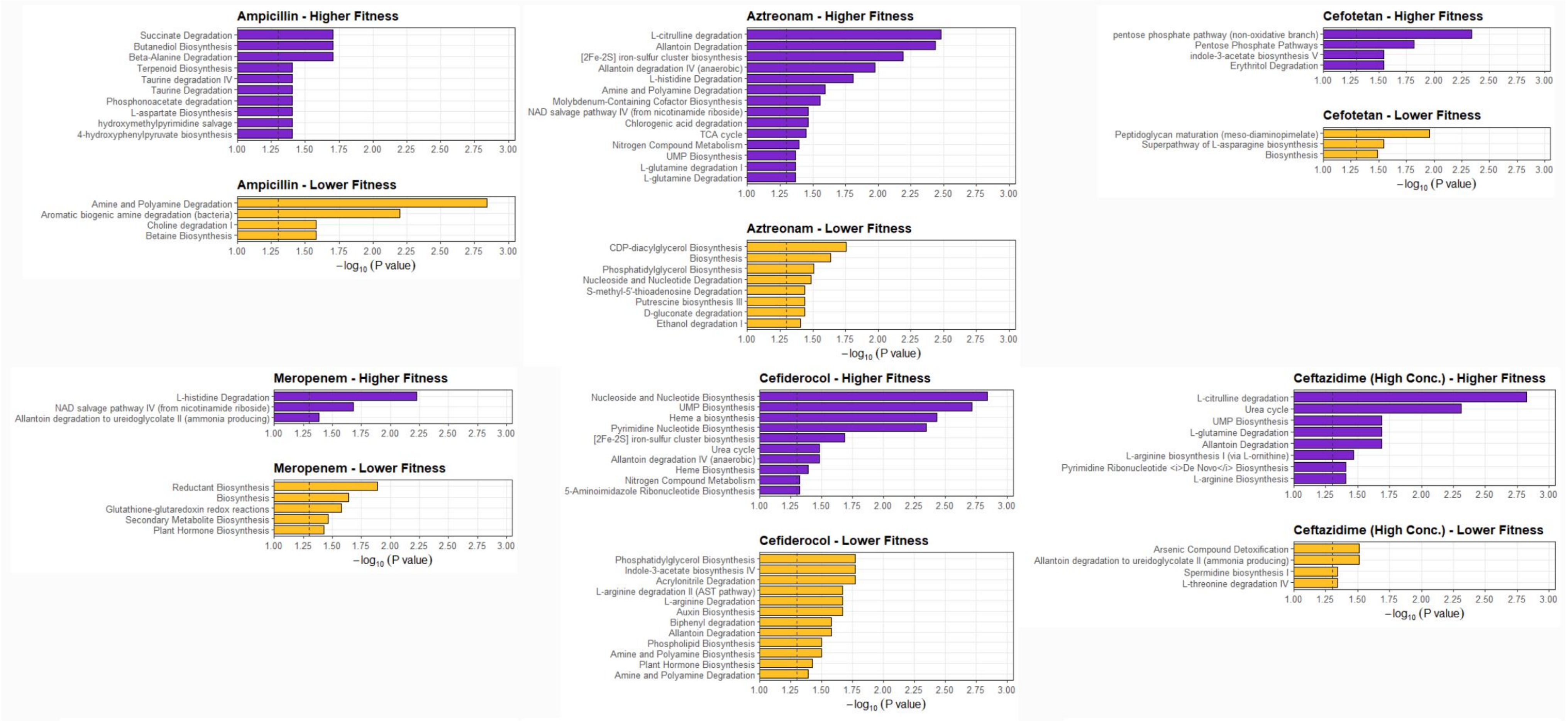

Ampicillin - Higher Fitness
Succinate Degradation
Butanediol Biosynthesis
Beta-Alanine Degradation
Terpenoid Biosynthesis
Taurine degradation IV
Taurine Degradation
Phosphonoacetate degradation
L-aspartate Biosynthesis
hydroxymethylpyrimidine salvage
4-hydroxyphenylpyruvate biosynthesis
1.00 1.25 1.50 1.75 2.00 2.25 2.50 2.75 3.00
Ampicillin - Lower Fitness
Amine and Polyamine Degradation
Aromatic biogenic amine degradation (bacteria)
Choline degradation I
Betaine Biosynthesis
1.00 1.25 1.50 1.75 2.00 2.25 2.50 2.75 3.00
$-\log_{10}$ (P value)
Aztreonam - Higher Fitness
L-citrulline degradation
Allantoin Degradation
[2Fe-2S] iron-sulfur cluster biosynthesis
Allantoin degradation IV (anaerobic)
L-histidine Degradation
Amine and Polyamine Degradation
Molybdenum-Containing Cofactor Biosynthesis
NAD salvage pathway IV (from nicotinamide riboside)
Chlorogenic acid degradation
TCA cycle
Nitrogen Compound Metabolism
UMP Biosynthesis
L-glutamine degradation I
L-glutamine Degradation
1.00 1.25 1.50 1.75 2.00 2.25 2.50 2.75 3.00
Aztreonam - Lower Fitness
CDP-diacylglycerol Biosynthesis
Biosynthesis
Phosphatidylglycerol Biosynthesis
Nucleoside and Nucleotide Degradation
S-methyl-5'-thioadenosine Degradation
Putrescine biosynthesis III
D-gluconate degradation
Ethanol degradation I
1.00 1.25 1.50 1.75 2.00 2.25 2.50 2.75 3.00
$-\log_{10}$ (P value)
Cefotetan - Higher Fitness
pentose phosphate pathway (non-oxidative branch)
Pentose Phosphate Pathways
indole-3-acetate biosynthesis V
Erythritol Degradation
1.00 1.25 1.50 1.75 2.00 2.25 2.50 2.75 3.00
Cefotetan - Lower Fitness
Peptidoglycan maturation (meso-diaminopimelate)
Superpathway of L-asparagine biosynthesis
Biosynthesis
1.00 1.25 1.50 1.75 2.00 2.25 2.50 2.75 3.00
$-\log_{10}$ (P value)
Meropenem - Higher Fitness
L-histidine Degradation
NAD salvage pathway IV (from nicotinamide riboside)
Allantoin degradation to ureidoglycolate II (ammonia producing)
1.00 1.25 1.50 1.75 2.00 2.25 2.50 2.75 3.00
Meropenem - Lower Fitness
Reductant Biosynthesis
Biosynthesis
Glutathione-glutaredoxin redox reactions
Secondary Metabolite Biosynthesis
Plant Hormone Biosynthesis
1.00 1.25 1.50 1.75 2.00 2.25 2.50 2.75 3.00
$-\log_{10}$ (P value)
Cefiderocol - Higher Fitness
Nucleoside and Nucleotide Biosynthesis
UMP Biosynthesis
Heme a biosynthesis
Pyrimidine Nucleotide Biosynthesis
[2Fe-2S] iron-sulfur cluster biosynthesis
Urea cycle
Allantoin degradation IV (anaerobic)
Heme Biosynthesis
Nitrogen Compound Metabolism
5-Aminoimidazole Ribonucleotide Biosynthesis
1.00 1.25 1.50 1.75 2.00 2.25 2.50 2.75 3.00
Cefiderocol - Lower Fitness
Phosphatidylglycerol Biosynthesis
Indole-3-acetate biosynthesis IV
Acrylonitrile Degradation
L-arginine degradation II (AST pathway)
L-arginine Degradation
Auxin Biosynthesis
Biphenyl degradation
Allantoin Degradation
Phospholipid Biosynthesis
Amine and Polyamine Biosynthesis
Plant Hormone Biosynthesis
Amine and Polyamine Degradation
1.00 1.25 1.50 1.75 2.00 2.25 2.50 2.75 3.00
$-\log_{10}$ (P value)
Ceftazidime (High Conc.) - Higher Fitness
L-citrulline degradation
Urea cycle
UMP Biosynthesis
L-glutamine Degradation
Allantoin Degradation
L-arginine biosynthesis I (via L-ornithine)
Pyrimidine Ribonucleotide <i>De Novo</i> Biosynthesis
L-arginine Biosynthesis
1.00 1.25 1.50 1.75 2.00 2.25 2.50 2.75 3.00
Ceftazidime (High Conc.) - Lower Fitness
Arsenic Compound Detoxification
Allantoin degradation to ureidoglycolate II (ammonia producing)
Spermidine biosynthesis I
L-threonine degradation IV
1.00 1.25 1.50 1.75 2.00 2.25 2.50 2.75 3.00
$-\log_{10}$ (P value)

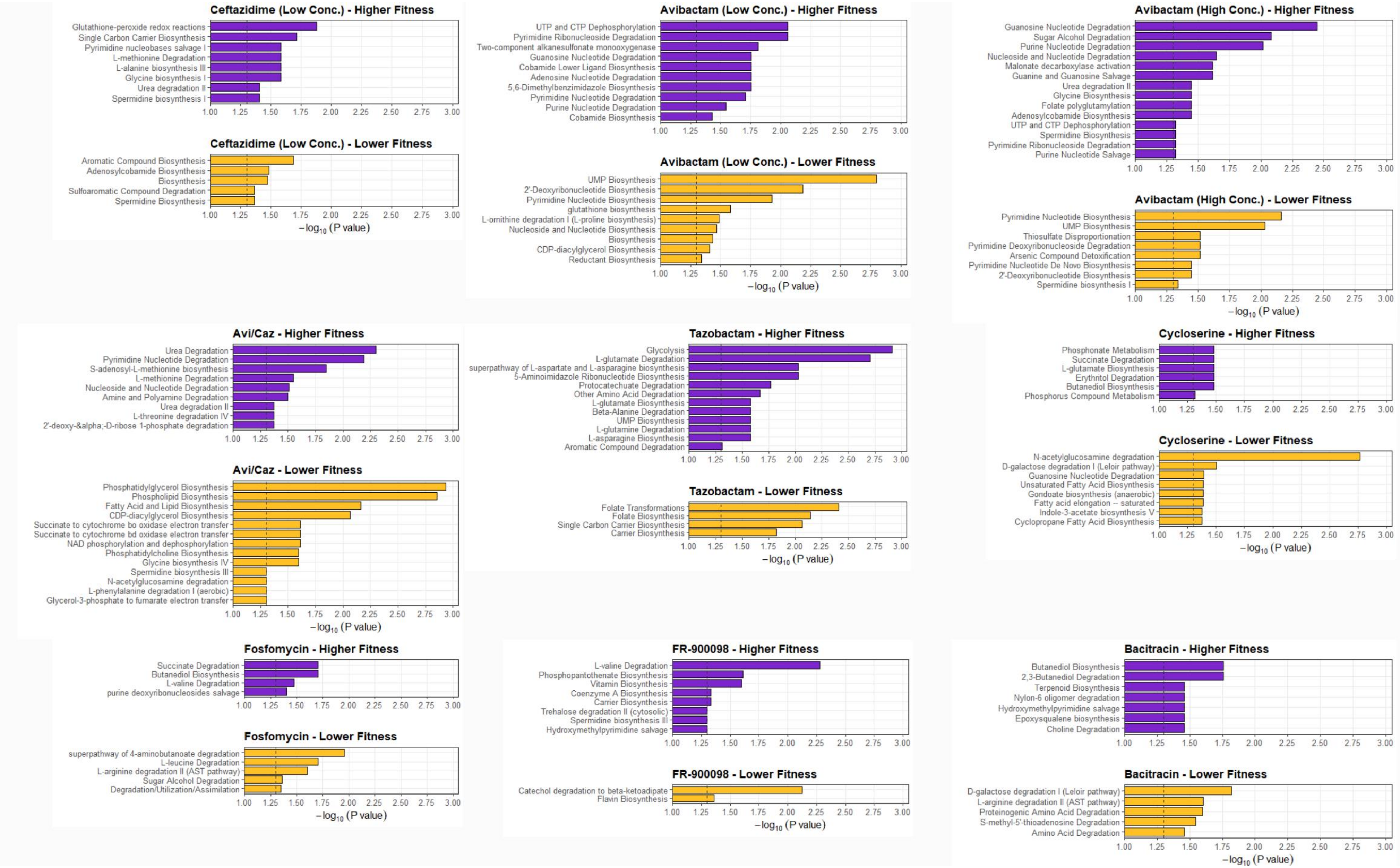
Ceftazidime (Low Conc.) - Higher Fitness
Glutathione-peroxide redox reactions
Single Carbon Carrier Biosynthesis
Pyrimidine nucleobases salvage I
L-methionine Degradation
L-alanine biosynthesis III
Glycine biosynthesis I
Urea degradation II
Spermidine biosynthesis I
Ceftazidime (Low Conc.) - Lower Fitness
Aromatic Compound Biosynthesis
Adenosylcobamide Biosynthesis
Biosynthesis
Sulfoaromatic Compound Degradation
Spermidine Biosynthesis
$-\log_{10}$ (P value)
Avibactam (Low Conc.) - Higher Fitness
UTP and CTP Dephosphorylation
Pyrimidine Ribonucleoside Degradation
Two-component alkanesulfonate monooxygenase
Guanosine Nucleotide Degradation
Cobamide Lower Ligand Biosynthesis
Adenosine Nucleotide Degradation
5,6-Dimethylbenzimidazole Biosynthesis
Pyrimidine Nucleotide Degradation
Purine Nucleotide Degradation
Cobamide Biosynthesis
Avibactam (Low Conc.) - Lower Fitness
UMP Biosynthesis
2'-Deoxyribonucleotide Biosynthesis
Pyrimidine Nucleotide Biosynthesis
glutathione biosynthesis
L-ornithine degradation I (L-proline biosynthesis)
Nucleoside and Nucleotide Biosynthesis
Biosynthesis
CDP-diacylglycerol Biosynthesis
Reductant Biosynthesis
$-\log_{10}$ (P value)
Avibactam (High Conc.) - Higher Fitness
Guanosine Nucleotide Degradation
Sugar Alcohol Degradation
Purine Nucleotide Degradation
Nucleoside and Nucleotide Degradation
Malonate decarboxylase activation
Guanine and Guanosine Salvage
Urea degradation II
Glycine Biosynthesis
Folate polyglutamylation
Adenosylcobamide Biosynthesis
UTP and CTP Dephosphorylation
Spermidine Biosynthesis
Pyrimidine Ribonucleoside Degradation
Purine Nucleotide Salvage
Avibactam (High Conc.) - Lower Fitness
Pyrimidine Nucleotide Biosynthesis
UMP Biosynthesis
Thiosulfate Disproportionation
Pyrimidine Deoxyribonucleoside Degradation
Arsenic Compound Detoxification
Pyrimidine Nucleotide De Novo Biosynthesis
2'-Deoxyribonucleotide Biosynthesis
Spermidine biosynthesis I
$-\log_{10}$ (P value)
Avi/Caz - Higher Fitness
Urea Degradation
Pyrimidine Nucleotide Degradation
S-adenosyl-L-methionine biosynthesis
L-methionine Degradation
Nucleoside and Nucleotide Degradation
Amine and Polyamine Degradation
Urea degradation II
L-threonine degradation IV
2'-deoxy-α-D-ribose 1-phosphate degradation
Avi/Caz - Lower Fitness
Phosphatidylglycerol Biosynthesis
Phospholipid Biosynthesis
Fatty Acid and Lipid Biosynthesis
CDP-diacylglycerol Biosynthesis
Succinate to cytochrome bo oxidase electron transfer
Succinate to cytochrome bd oxidase electron transfer
NAD phosphorylation and dephosphorylation
Phosphatidylcholine Biosynthesis
Glycine biosynthesis IV
Spermidine biosynthesis III
N-acetylglucosamine degradation
L-phenylalanine degradation I (aerobic)
Glycerol-3-phosphate to fumarate electron transfer
$-\log_{10}$ (P value)
Tazobactam - Higher Fitness
Glycolysis
L-glutamate Degradation
superpathway of L-aspartate and L-asparagine biosynthesis
5-Aminoimidazole Ribonucleotide Biosynthesis
Protocatechuate Degradation
Other Amino Acid Degradation
L-glutamate Biosynthesis
Beta-Alanine Degradation
UMP Biosynthesis
L-glutamine Degradation
L-asparagine Biosynthesis
Aromatic Compound Degradation
Tazobactam - Lower Fitness
Folate Transformations
Folate Biosynthesis
Single Carbon Carrier Biosynthesis
Carrier Biosynthesis
$-\log_{10}$ (P value)
Cycloserine - Higher Fitness
Phosphonate Metabolism
Succinate Degradation
L-glutamate Biosynthesis
Erythritol Degradation
Butanediol Biosynthesis
Phosphorus Compound Metabolism
Cycloserine - Lower Fitness
N-acetylglucosamine degradation
D-galactose degradation I (Leloir pathway)
Guanosine Nucleotide Degradation
Unsaturated Fatty Acid Biosynthesis
Gondoate biosynthesis (anaerobic)
Fatty acid elongation -- saturated
Indole-3-acetate biosynthesis V
Cyclopropane Fatty Acid Biosynthesis
$-\log_{10}$ (P value)
Fosfomycin - Higher Fitness
Succinate Degradation
Butanediol Biosynthesis
L-valine Degradation
purine deoxyribonucleosides salvage
Fosfomycin - Lower Fitness
superpathway of 4-aminobutanoate degradation
L-leucine Degradation
L-arginine degradation II (AST pathway)
Sugar Alcohol Degradation
Degradation/Utilization/Assimilation
$-\log_{10}$ (P value)
FR-900098 - Higher Fitness
L-valine Degradation
Phosphopantothenate Biosynthesis
Vitamin Biosynthesis
Coenzyme A Biosynthesis
Carrier Biosynthesis
Trehalose degradation II (cytosolic)
Spermidine biosynthesis III
Hydroxymethylpyrimidine salvage
FR-900098 - Lower Fitness
Catechol degradation to beta-ketoadipate
Flavin Biosynthesis
$-\log_{10}$ (P value)
Bacitracin - Higher Fitness
Butanediol Biosynthesis
2,3-Butanediol Degradation
Terpenoid Biosynthesis
Nylon-6 oligomer degradation
Hydroxymethylpyrimidine salvage
Epoxysqualene biosynthesis
Choline Degradation
Bacitracin - Lower Fitness
D-galactose degradation I (Leloir pathway)
L-arginine degradation II (AST pathway)
Proteinogenic Amino Acid Degradation
S-methyl-5'-thioadenosine Degradation
Amino Acid Degradation
1.00 1.25 1.50 1.75 2.00 2.25 2.50 2.75 3.00
$-\log_{10}$ (P value)

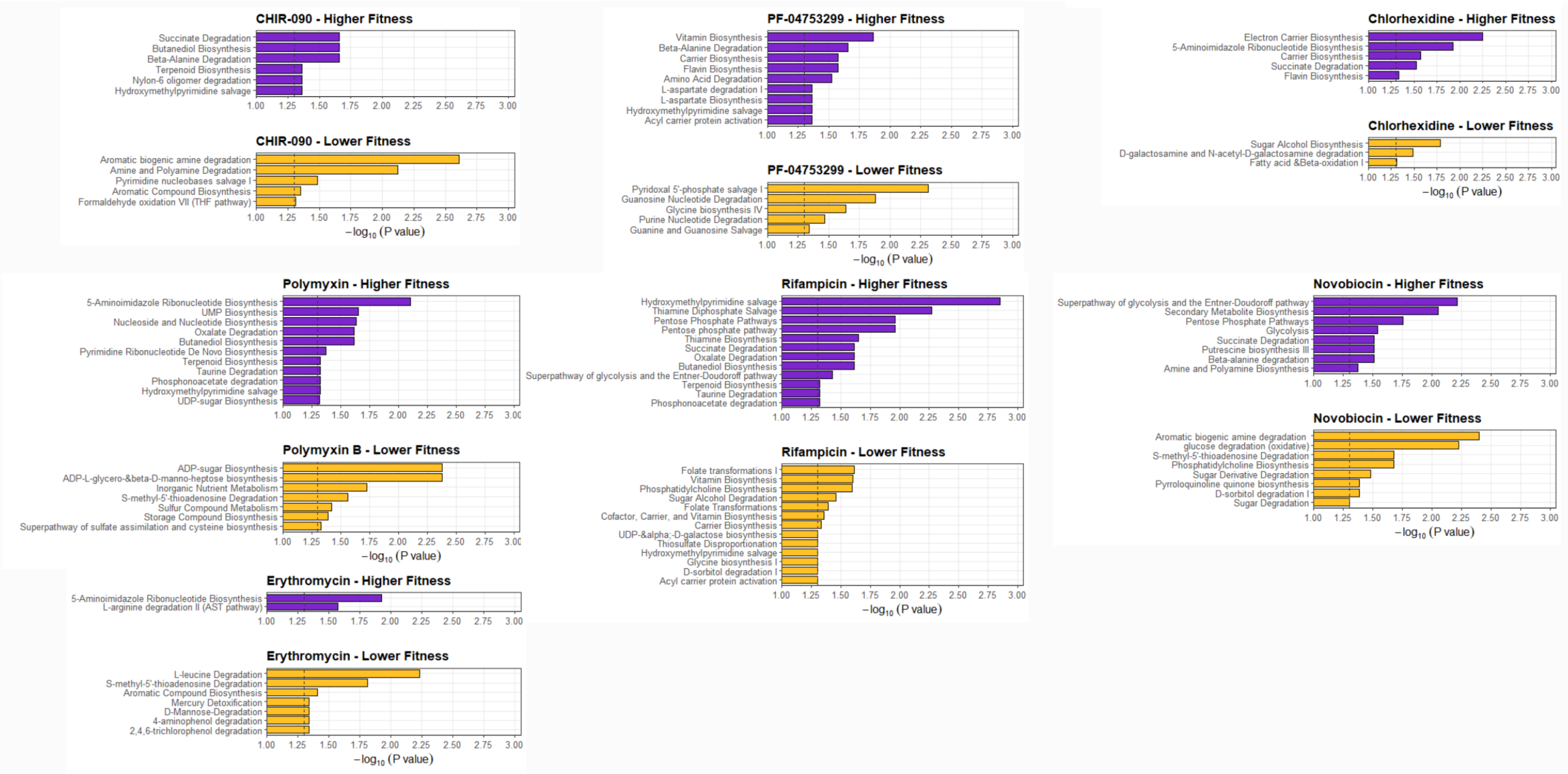


**Figure S6. BioCyc pathway enrichment of genes affecting fitness in the presence of the antibiotic panel.** Genes with fitness scores either above 0.5 or below -0.5 were submitted to BioCyc for enrichment analysis (Fisher exact test without multiple testing correction) (against K56-2 genome from GenBank accession GCA_000333155.2). Enrichments from genes with positive and negative fitness scores are given in purple and gold, respectively. The dashed line indicates P = 0.05.

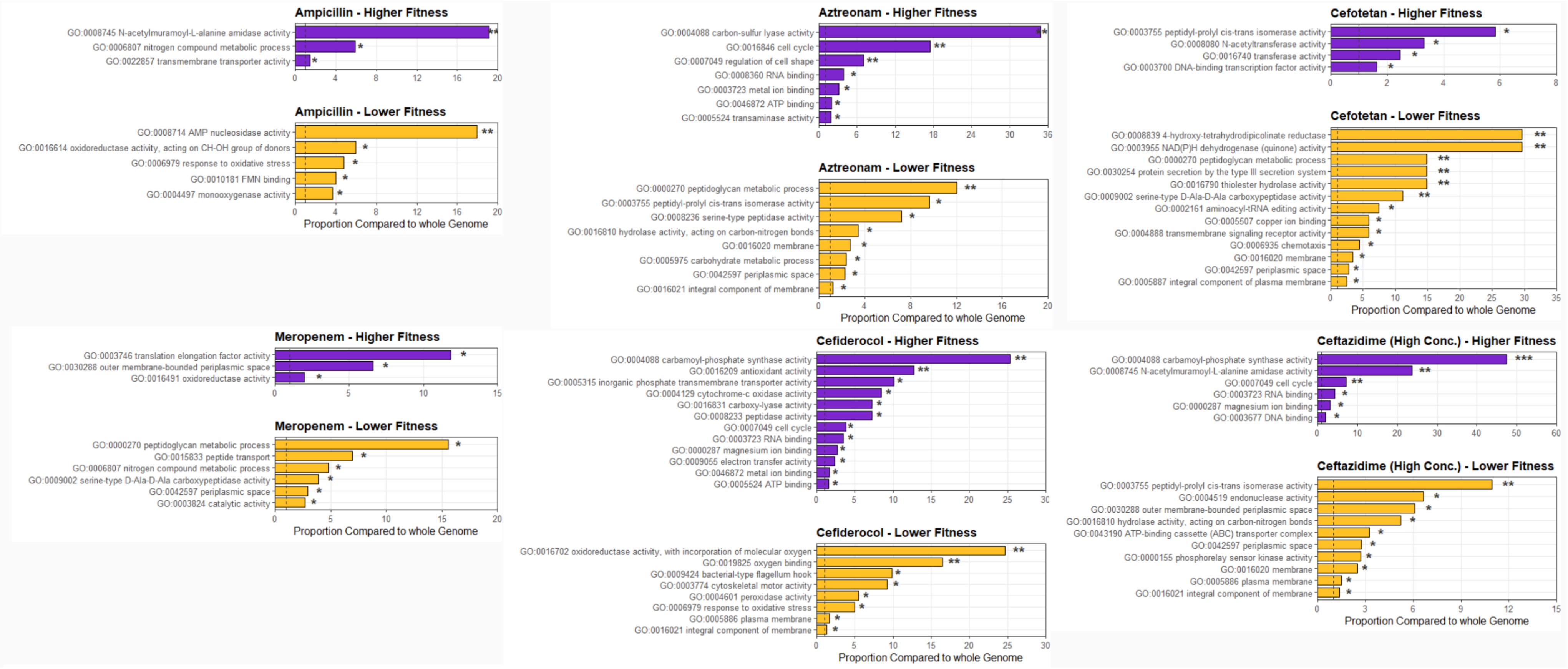

Ampicillin - Higher Fitness
GO:0008745 N-acetylmuramoyl-L-alanine amidase activity
GO:0006807 nitrogen compound metabolic process
GO:0022857 transmembrane transporter activity
Ampicillin - Lower Fitness
GO:0008714 AMP nucleosidase activity
GO:0016614 oxidoreductase activity, acting on CH-OH group of donors
GO:0006979 response to oxidative stress
GO:0010181 FMN binding
GO:0004497 monooxygenase activity
Proportion Compared to whole Genome
Aztreonam - Higher Fitness
GO:0004088 carbon-sulfur lyase activity
GO:0016846 cell cycle
GO:0007049 regulation of cell shape
GO:0008360 RNA binding
GO:0003723 metal ion binding
GO:0046872 ATP binding
GO:0005524 transaminase activity
Aztreonam - Lower Fitness
GO:0000270 peptidoglycan metabolic process
GO:0003755 peptidyl-prolyl cis-trans isomerase activity
GO:0008236 serine-type peptidase activity
GO:0016810 hydrolase activity, acting on carbon-nitrogen bonds
GO:0016020 membrane
GO:0005975 carbohydrate metabolic process
GO:0042597 periplasmic space
GO:0016021 integral component of membrane
Proportion Compared to whole Genome
Cefotetan - Higher Fitness
GO:0003755 peptidyl-prolyl cis-trans isomerase activity
GO:0008080 N-acetyltransferase activity
GO:0016740 transferase activity
GO:0003700 DNA-binding transcription factor activity
Cefotetan - Lower Fitness
GO:0008839 4-hydroxy-tetrahydrodipicolinate reductase
GO:0003955 NAD(P)H dehydrogenase (quinone) activity
GO:0000270 peptidoglycan metabolic process
GO:0030254 protein secretion by the type III secretion system
GO:0016790 thiolester hydrolase activity
GO:0009002 serine-type D-Ala-D-Ala carboxypeptidase activity
GO:0002161 aminoacyl-tRNA editing activity
GO:0005507 copper ion binding
GO:0004888 transmembrane signaling receptor activity
GO:0006935 chemotaxis
GO:0016020 membrane
GO:0042597 periplasmic space
GO:0005887 integral component of plasma membrane
Proportion Compared to whole Genome
Meropenem - Higher Fitness
GO:0003746 translation elongation factor activity
GO:0030288 outer membrane-bounded periplasmic space
GO:0016491 oxidoreductase activity
Meropenem - Lower Fitness
GO:0000270 peptidoglycan metabolic process
GO:0015833 peptide transport
GO:0006807 nitrogen compound metabolic process
GO:0009002 serine-type D-Ala-D-Ala carboxypeptidase activity
GO:0042597 periplasmic space
GO:0003824 catalytic activity
Proportion Compared to whole Genome
Cefiderocol - Higher Fitness
GO:0004088 carbamoyl-phosphate synthase activity
GO:0016209 antioxidant activity
GO:0005315 inorganic phosphate transmembrane transporter activity
GO:0004129 cytochrome-c oxidase activity
GO:0016831 carboxy-lyase activity
GO:0008233 peptidase activity
GO:0007049 cell cycle
GO:0003723 RNA binding
GO:0000287 magnesium ion binding
GO:0009055 electron transfer activity
GO:0046872 metal ion binding
GO:0005524 ATP binding
Cefiderocol - Lower Fitness
GO:0016702 oxidoreductase activity, with incorporation of molecular oxygen
GO:0019825 oxygen binding
GO:0009424 bacterial-type flagellum hook
GO:0003774 cytoskeletal motor activity
GO:0004601 peroxidase activity
GO:0006979 response to oxidative stress
GO:0005886 plasma membrane
GO:0016021 integral component of membrane
Proportion Compared to whole Genome
Ceftazidime (High Conc.) - Higher Fitness
GO:0004088 carbamoyl-phosphate synthase activity
GO:0008745 N-acetylmuramoyl-L-alanine amidase activity
GO:0007049 cell cycle
GO:0003723 RNA binding
GO:0000287 magnesium ion binding
GO:0003677 DNA binding
Ceftazidime (High Conc.) - Lower Fitness
GO:0003755 peptidyl-prolyl cis-trans isomerase activity
GO:0004519 endonuclease activity
GO:0030288 outer membrane-bounded periplasmic space
GO:0016810 hydrolase activity, acting on carbon-nitrogen bonds
GO:0043190 ATP-binding cassette (ABC) transporter complex
GO:0042597 periplasmic space
GO:0000155 phosphorelay sensor kinase activity
GO:0016020 membrane
GO:0005886 plasma membrane
GO:0016021 integral component of membrane
Proportion Compared to whole Genome

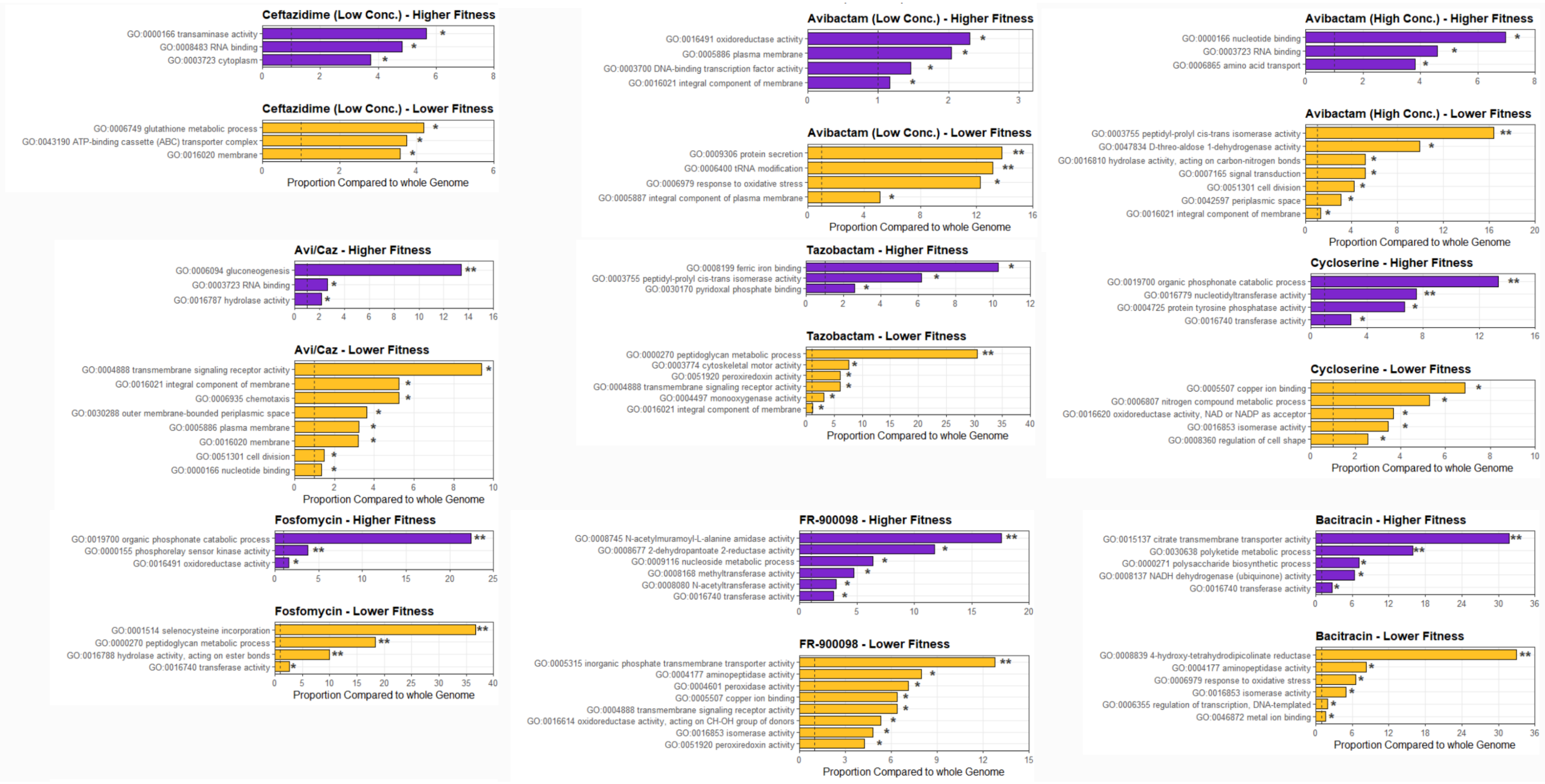
Ceftazidime (Low Conc.) - Higher Fitness
GO:0000166 transaminase activity
GO:0008483 RNA binding
GO:0003723 cytoplasm
Ceftazidime (Low Conc.) - Lower Fitness
GO:0006749 glutathione metabolic process
GO:0043190 ATP-binding cassette (ABC) transporter complex
GO:0016020 membrane
Proportion Compared to whole Genome
Avibactam (Low Conc.) - Higher Fitness
GO:0016491 oxidoreductase activity
GO:0005886 plasma membrane
GO:0003700 DNA-binding transcription factor activity
GO:0016021 integral component of membrane
Avibactam (Low Conc.) - Lower Fitness
GO:0009306 protein secretion
GO:0006400 tRNA modification
GO:0006979 response to oxidative stress
GO:0005887 integral component of plasma membrane
Proportion Compared to whole Genome
Avibactam (High Conc.) - Higher Fitness
GO:0000166 nucleotide binding
GO:0003723 RNA binding
GO:0006865 amino acid transport
Avibactam (High Conc.) - Lower Fitness
GO:0003755 peptidyl-prolyl cis-trans isomerase activity
GO:0047834 D-threo-aldose 1-dehydrogenase activity
GO:0016810 hydrolase activity, acting on carbon-nitrogen bonds
GO:0007165 signal transduction
GO:0051301 cell division
GO:0042597 periplasmic space
GO:0016021 integral component of membrane
Proportion Compared to whole Genome
Avi/Caz - Higher Fitness
GO:0006094 gluconeogenesis
GO:0003723 RNA binding
GO:0016787 hydrolase activity
Avi/Caz - Lower Fitness
GO:0004888 transmembrane signaling receptor activity
GO:0016021 integral component of membrane
GO:0006935 chemotaxis
GO:0030288 outer membrane-bounded periplasmic space
GO:0005886 plasma membrane
GO:0016020 membrane
GO:0051301 cell division
GO:0000166 nucleotide binding
Proportion Compared to whole Genome
Tazobactam - Higher Fitness
GO:0008199 ferric iron binding
GO:0003755 peptidyl-prolyl cis-trans isomerase activity
GO:0030170 pyridoxal phosphate binding
Tazobactam - Lower Fitness
GO:0000270 peptidoglycan metabolic process
GO:0003774 cytoskeletal motor activity
GO:0051920 peroxiredoxin activity
GO:0004888 transmembrane signaling receptor activity
GO:0004497 monooxygenase activity
GO:0016021 integral component of membrane
Proportion Compared to whole Genome
Cycloserine - Higher Fitness
GO:0019700 organic phosphonate catabolic process
GO:0016779 nucleotidyltransferase activity
GO:0004725 protein tyrosine phosphatase activity
GO:0016740 transferase activity
Cycloserine - Lower Fitness
GO:0005507 copper ion binding
GO:0006807 nitrogen compound metabolic process
GO:0016620 oxidoreductase activity, NAD or NADP as acceptor
GO:0016853 isomerase activity
GO:0008360 regulation of cell shape
Proportion Compared to whole Genome
Fosfomycin - Higher Fitness
GO:0019700 organic phosphonate catabolic process
GO:0000155 phosphorelay sensor kinase activity
GO:0016491 oxidoreductase activity
Fosfomycin - Lower Fitness
GO:0001514 selenocysteine incorporation
GO:0000270 peptidoglycan metabolic process
GO:0016788 hydrolase activity, acting on ester bonds
GO:0016740 transferase activity
Proportion Compared to whole Genome
FR-900098 - Higher Fitness
GO:0008745 N-acetylmuramoyl-L-alanine amidase activity
GO:0008677 2-dehydropantoate 2-reductase activity
GO:0009116 nucleoside metabolic process
GO:0008168 methyltransferase activity
GO:0008080 N-acetyltransferase activity
GO:0016740 transferase activity
FR-900098 - Lower Fitness
GO:0005315 inorganic phosphate transmembrane transporter activity
GO:0004177 aminopeptidase activity
GO:0004601 peroxidase activity
GO:0005507 copper ion binding
GO:0004888 transmembrane signaling receptor activity
GO:0016614 oxidoreductase activity, acting on CH-OH group of donors
GO:0016853 isomerase activity
GO:0051920 peroxiredoxin activity
Proportion Compared to whole Genome
Bacitracin - Higher Fitness
GO:0015137 citrate transmembrane transporter activity
GO:0030638 polyketide metabolic process
GO:0000271 polysaccharide biosynthetic process
GO:0008137 NADH dehydrogenase (ubiquinone) activity
GO:0016740 transferase activity
Bacitracin - Lower Fitness
GO:0008839 4-hydroxy-tetrahydrodipicolinate reductase
GO:0004177 aminopeptidase activity
GO:0006979 response to oxidative stress
GO:0016853 isomerase activity
GO:0006355 regulation of transcription, DNA-templated
GO:0046872 metal ion binding
Proportion Compared to whole Genome

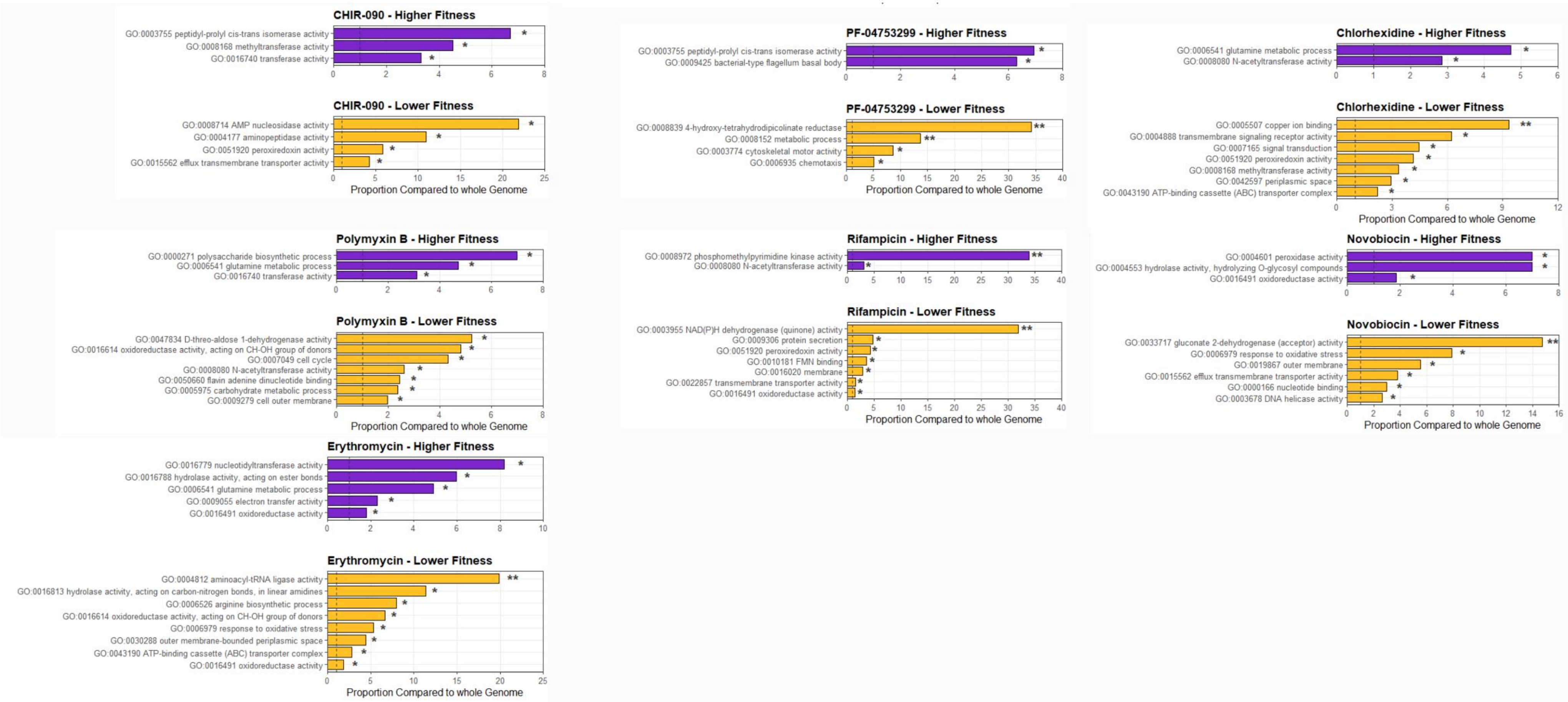


**Figure S7.** GO term enrichment of genes affecting fitness in the presence of the antibiotic panel. Genes with fitness scores either above 0.5 or below -0.5 were used for GO term enrichment relative to the whole K56-2 genome (Hypergeometric test without multiple testing correction) with GeneMerge 1.5 (Castillo-Davis et al. 2003). Enrichments from genes with positive and negative fitness scores are given in purple and gold, respectively. The dashed line indicates equal proportions in the genome and antibiotic condition. * P < 0.05, ** P < 0.01, *** P < 0.001.

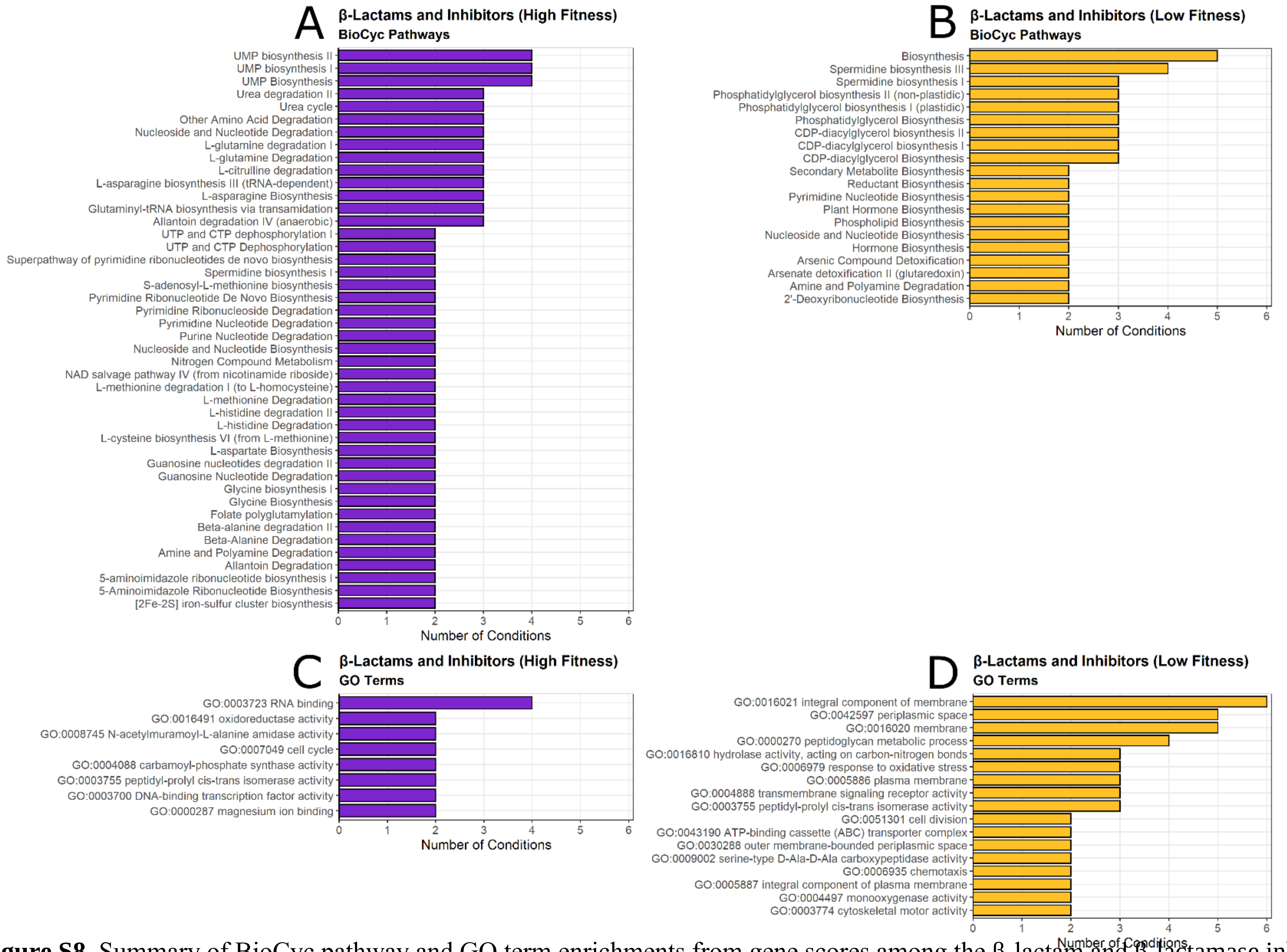


**Figure S8.** Summary of BioCyc pathway and GO term enrichments from gene scores among the β-lactam and β-lactamase inhibitor conditions. Significant enrichments ($P < 0.05$) from BioCyc pathways and GO terms were pooled from Figures A4 and A5. Only those pathways and terms present in at least two antibiotic conditions are shown.

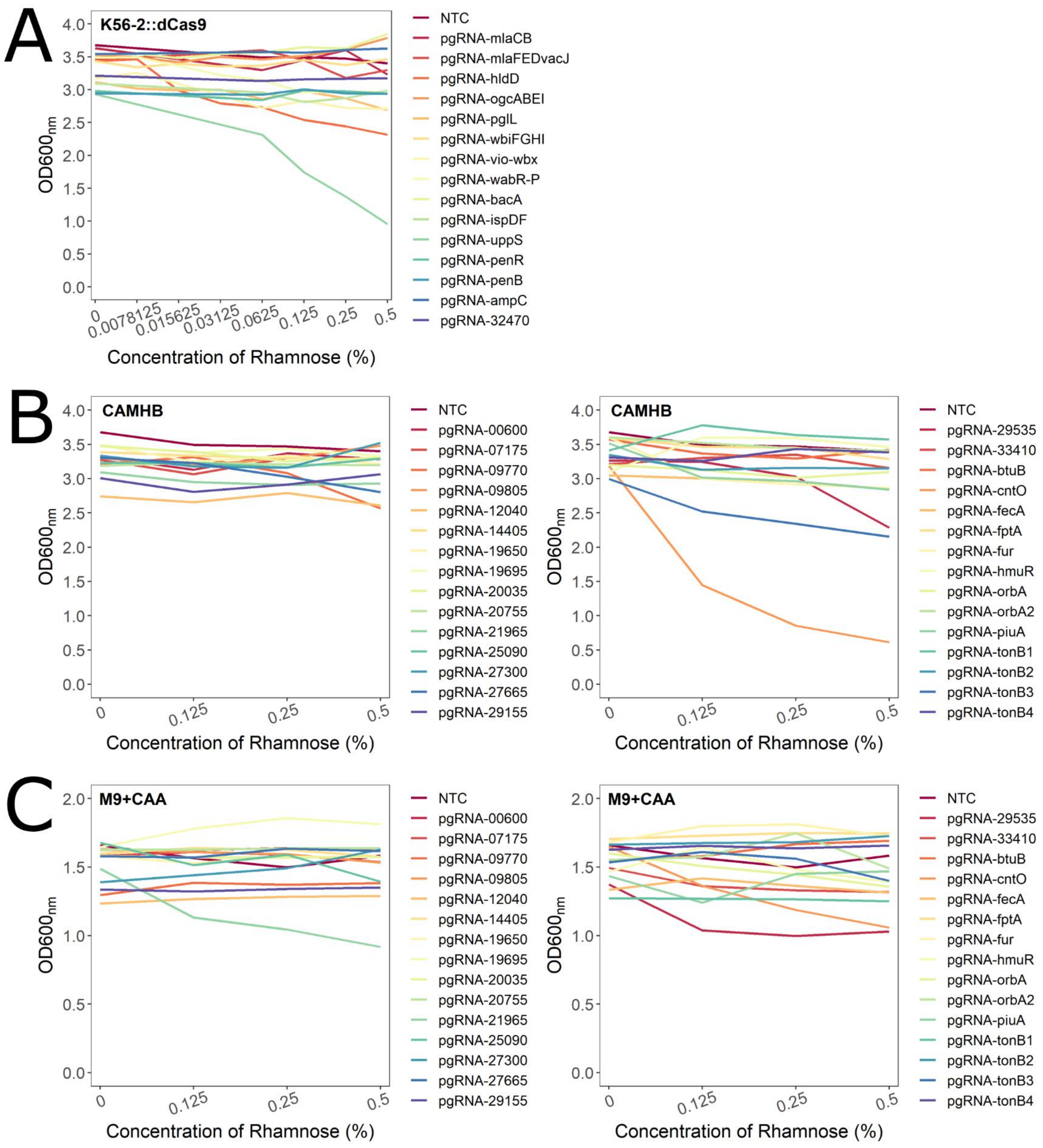
A
K56-2::dCas9
OD600nm
Concentration of Rhamnose (%)
0
0.0078125
0.015625
0.03125
0.0625
0.125
0.25
0.5
NTC
pgRNA-mlaCB
pgRNA-mlaFEDvacJ
pgRNA-hldD
pgRNA-ogcABEI
pgRNA-pglL
pgRNA-wbiFGHI
pgRNA-vio-wbx
pgRNA-wabR-P
pgRNA-bacA
pgRNA-ispDF
pgRNA-uppS
pgRNA-penR
pgRNA-penB
pgRNA-ampC
pgRNA-32470
B
CAMHB
NTC
pgRNA-00600
pgRNA-07175
pgRNA-09770
pgRNA-09805
pgRNA-12040
pgRNA-14405
pgRNA-19650
pgRNA-19695
pgRNA-20035
pgRNA-20755
pgRNA-21965
pgRNA-25090
pgRNA-27300
pgRNA-27665
pgRNA-29155
NTC
pgRNA-29535
pgRNA-33410
pgRNA-btuB
pgRNA-cntO
pgRNA-fecA
pgRNA-fptA
pgRNA-fur
pgRNA-hmuR
pgRNA-orbA
pgRNA-orbA2
pgRNA-piuA
pgRNA-tonB1
pgRNA-tonB2
pgRNA-tonB3
pgRNA-tonB4
C
M9+CAA

**Figure S9.** Rhamnose dose-responses of select CRISPRi mutants constructed in this study. Cultures were grown with the indicated concentrations of rhamnose in A) LB, B) CAMHB, C) M9+CAA for 16 hours, the time at which the control strains reached the maximum $OD_{600}$, and then $OD_{600}$ values were recorded Each gene was targeted with two sgRNA, and one representative is shown here. If the targeted gene does not have a 4-letter name, the locus tag (preceded by K562_) is given. NTC = non-targeting control sgRNA. Values shown are averages of at least three biological replicates; SD values are omitted for clarity.

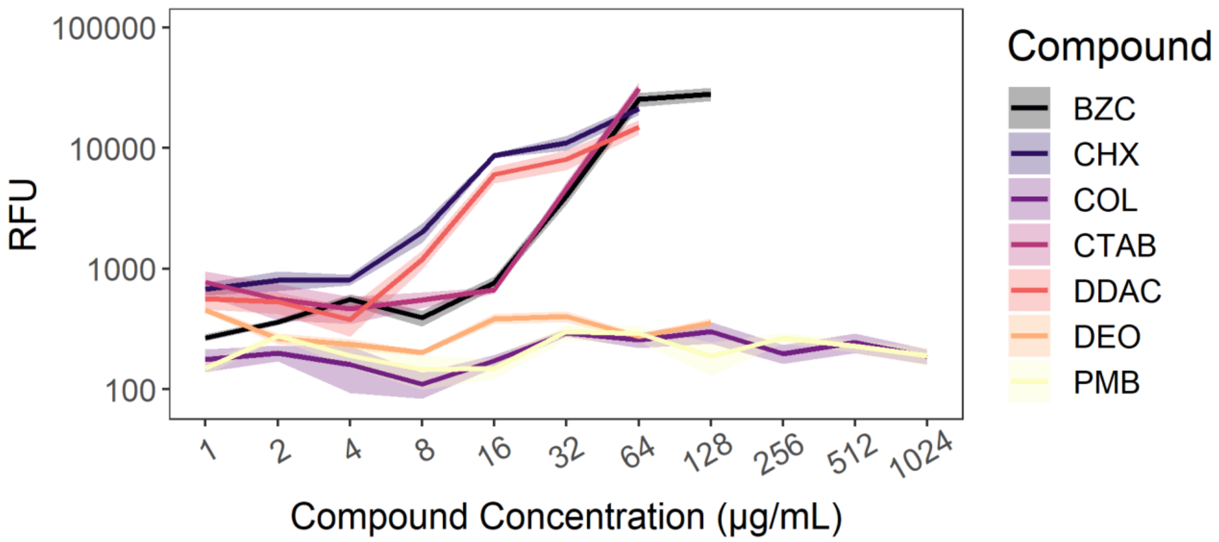


**Figure S10.** Outer membrane permeability of K56-2 is increased by exposure to certain membrane disruptors. Exponential phase K56-2 was incubated with NPN and a concentration gradient of the indicated compounds. BZC = benzethonium chloride; CHX = chlorhexidine; COL = colistin; CTAB = cetrimonium bromide; DDAC = dodecyl dimethylammonium bromide; DEO = sodium deoxycholate; PMB = polymyxin B. Raw fluorescence values were blank corrected. The shading represents means ± SD of three biological replicates.

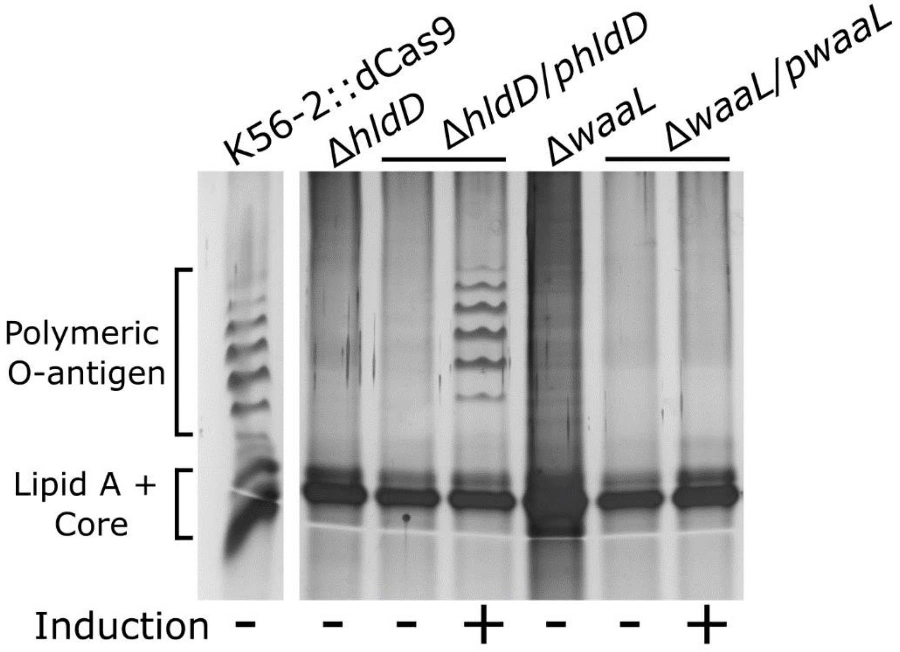


**Figure S11**. Effect of deletion and complementation of *hldD* and *waaL* on O-antigen expression. These genes were deleted in the K56-2::dCas9 background. Silver-stained SDS polyacrylamide gels of LPS extracts from mid-exponential phase deletion and complementation mutants grown for 4 hours with or without 0.05% rhamnose. Shown are different lanes from the same gel.

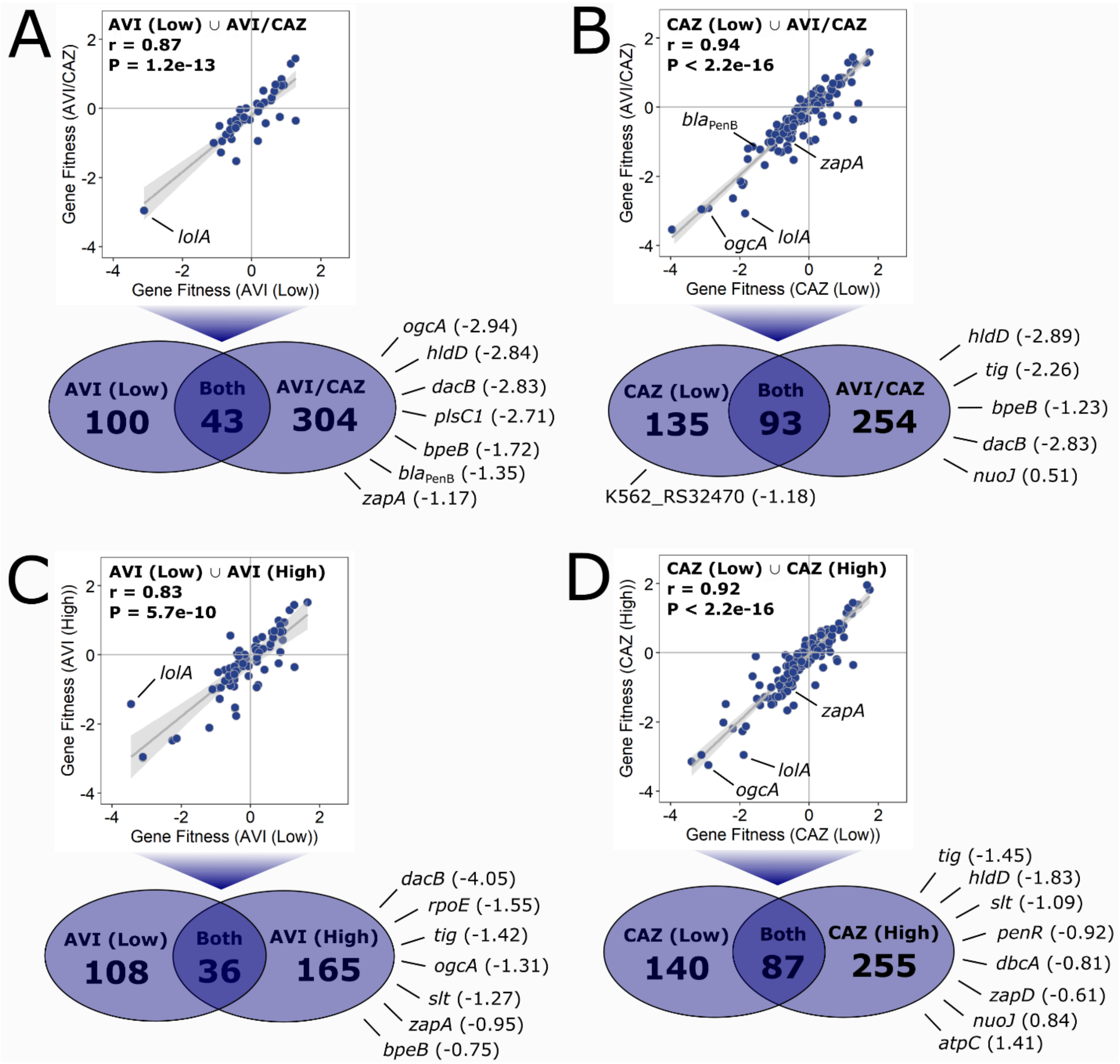


**Figure S12.** Concentration- and combination-dependent chemical-genetic interactions with AVI/CAZ. Comparison gene fitness scores between A) AVI/CAZ to AVI-L, B) AVI/CAZ to CAZ-L, C) AVI-H to AVI-L, and D) CAZ-H to CAZ-L. All genes with significantly different fitness scores greater than 0.5 or less than -0.5 relative to the DMSO control ($P < 0.05$) were used for comparison. For the genes that were common to both conditions, the scatter plots show a Pearson's correlation test (with 95% confidence interval in light grey) and associated r and P-values. Fitness scores next to genes are relative to the DMSO control.

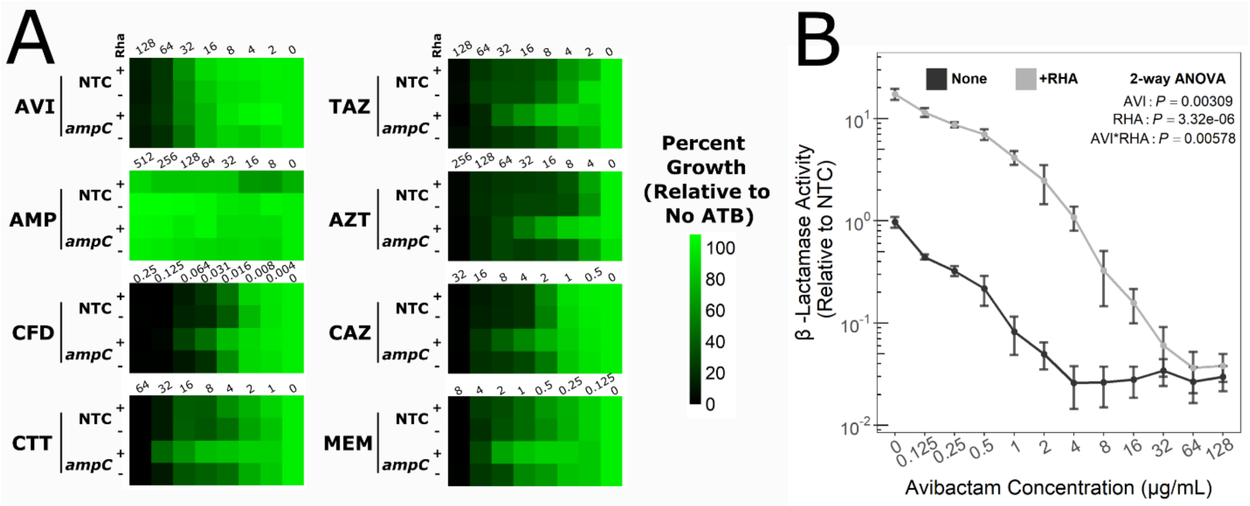


**Figure S13.** Knockdown of *bla*$_{AmpC}$ reduces β-lactam susceptibility but is still rescued by AVI. A) Antibiotic dose responses (µg/mL) of growth of the indicated mutants with or without CRISPRi induction with 0.5% rhamnose. Values are normalized to the $OD_{600}$ of growth without antibiotic, and are means of three biologica replicates. NTC = non-targeting sgRNA control. B) Nitrocefin hydrolysis assay of the knockdown mutant in *bla*$_{AmpC}$ incubated with increasing concentrations of AVI with or without CRISPRi induction. Data was normalized to the NTC mutant. Values presented are means of four biological replicates, ± SD. Significance was determined by a two-way ANOVA.
